# Membrane Fusion Inhibition, Immune Modulation, and Cholesterol Synthesis Dysregulation During Dengue Virus Inhibition by 25-Hydroxycholesterol

**DOI:** 10.1101/2025.05.31.657153

**Authors:** Debayani Chakraborty, Achinta Sannigrahi, Mahammed Kaif, Arijit Patra, Deep Thakkar, Debarati Chakraborty, Rahul Roy

## Abstract

Physicochemical properties and composition of cellular membranes are crucial for regulating broad cellular responses including signaling and defense against pathogens. Dengue virus (DENV) exploits cholesterol-rich membranes and host lipid pathways, such as cholesterol biosynthesis, lipid raft organization, and lipid droplet formation, for entry, replication, and assembly. Additionally, lipid-based plasma membrane signaling can trigger innate immune responses that attenuate viral growth, underscoring the dual role of lipids in facilitating and restricting DENV infection. Here, we demonstrate that 25-hydroxycholesterol (25-HC), an oxidized cholesterol metabolite, inhibits DENV infection through a multifaceted mechanism. 25-HC disrupts viral membrane fusion by altering cholesterol distribution and lipid raft organization, impairing the binding and fusion of the DENV envelope (E) protein with host membranes. Additionally, 25-HC modulates host cholesterol metabolism by suppressing biosynthesis pathways essential for viral replication while enhancing lipid droplet formation and stress-response pathways. Transcriptomic analyses reveal that 25-HC primes innate immune responses, activating proinflammatory pathways such as the NLRP3 inflammasome and MAPK signaling, while selectively modulating interferon-stimulated gene expression. Notably, 25-HC exhibits synergistic antiviral effects when combined with direct-acting antivirals like Remdesivir, underscoring its potential in combination therapies. These findings establish 25-HC as a promising candidate for host-directed antiviral strategies against DENV and other enveloped viruses.

## 1 INTRODUCTION

The cell membrane is a dynamic and responsive structure, constantly adapting to physiological and pathological stimuli to maintain homeostasis. This adaptability becomes particularly critical during pathogen invasion, where the membrane serves as the first point of contact, enabling the activation of signaling pathways and modifications to its composition to counteract infection[1, 2, 3]. Enveloped viruses, for instance, exploit this dynamic nature through viral membrane fusion—a critical step that allows them to deliver genetic material into the host cells and initiate infection[2, 4]. This process is mediated by viral glycoproteins, which undergo conformational changes to merge the viral envelope with the host cell membrane[5]. Viruses such as flaviviruses, coronaviruses, and filoviruses depend on these glycoproteins to interact with host receptors and trigger fusion at the plasma membrane or within endosomal compartments[4, 6, 7, 8, 9, 10]. Beyond viral glycoproteins, host cell membrane lipids, such as cholesterol and sphingolipids, play a pivotal role in modulating the efficiency and dynamics of viral fusion by influencing membrane curvature and fluidity [11, 8, 12, 2].

Given the critical role of cholesterol in membrane function and homeostasis, it is not surprising that many viruses exploit cholesterol-rich microdomains to facilitate membrane fusion and infection[13, 14, 11]. These cholesterol-dependent processes make the host membrane an attractive target for antiviral strategies. Among the potential membrane targeting approaches, oxysterols like 25-hydroxycholesterol (25-HC) have emerged as promising antivirals[15, 16]. Produced by the enzyme cholesterol 25-hydroxylase (CH25H) in response to interferon signaling, 25-HC disrupts viral entry by altering the lipid composition of host membranes and disrupting the entry process[17, 15, 18]. Its broad-spectrum activity against enveloped viruses [19, 18, 20, 21] and even non-enveloped viruses like human Rotavirus [22] have raised interest in using such membrane-active oxysterols to disrupt viral entry in inhibiting infection.

Beyond its direct antiviral action by modulating cell membrane properties, 25-HC also acts as an immunomodulator. It regulates inflammatory cytokines, such as interleukin-6 (IL-6) and tumor necrosis factor-alpha (TNF-α), by inhibiting nuclear factor kappa B (NF-κB) activation [23, 24]. This helps prevent excessive inflammation and tissue damage, maintaining immune homeostasis [25]. Furthermore, 25-HC modulates type I interferon (IFN) responses, enhancing antiviral defences while limiting immune overactivation [17, 26]. A critical aspect of 25-HC’s immunomodulatory role is its regulation of cholesterol metabolism, which is tightly linked to immune function. Cellular cholesterol is distributed across different pools, such as free and sphingomyelin-bound cholesterol in the plasma membrane and ER. Cholesterol synthesis is regulated by ER transmembrane proteins, including SCAP and SREBP, which respond to cholesterol availability [27, 28]. By suppressing SREBP activity, 25-HC reduces cholesterol synthesis and uptake, limiting the cholesterol required for viral replication and assembly [29, 30].

Despite significant progress in understanding the antiviral properties of 25-HC, key gaps remain in elucidating its mechanisms of action. While 25-HC disrupts viral entry at the host plasma membrane, the precise molecular details—such as its effects on membrane properties and organization—leading to entry inhibition are not fully understood. It is unclear whether 25-HC directly interacts with viral glycoproteins, host receptors, or if its effects are entirely mediated through changes in the membrane. Furthermore, the extent to which its immunomodulatory effects, including modulation of type I interferon signaling, inflammatory cytokine expression, and activation of pathways like NLRP3 inflammasome and MAPK signaling, contribute to its antiviral activity remains poorly defined. Key questions include whether these immune-modulating properties act synergistically with its membrane-targeting effects or function independently, and whether 25-HC’s suppression of cholesterol biosynthesis inadvertently promotes lipid droplet formation, which could support viral replication.

Although 25-HC has been examined with regard to various viruses, its precise function in altering Dengue virus (DENV) infection is not known. Critically, it remains unclear whether 25-HC impairs the binding and fusion processes of the DENV envelope (E) protein with host cell membranes, changes cholesterol distribution essential for viral entry, or affects additional phases of the viral life cycle, including replication, assembly, or modulation of the host immune response. Moreover, a comprehensive understanding of how 25-HC could inhibit or alter specific immune responses, especially when used in combination with antiviral therapies, is essential for boosting its synergistic effects and guiding the development of more potent antiviral strategies. While it could potentially increase the effectiveness of certain antiviral agents, the mechanisms responsible for these effects remain unclear, and identifying the most efficient combinations might aid in the development of robust therapeutic strategies.

In this study, we investigate the underlying mechanism of inihibition of Dengue virus 2 (DENV2) by 25-HC. We demonstrate that 25-HC disrupts DENV2 infection by membrane fusion inhibition as well as induces cholesterol biosynthesis dysregulation and pro-inflammatory immune response. We elucidate this 25-HC action during DENV infection using a combination of *in vitro* and *in cellulo* fusion assays, time-of-addition virus infection assays, and transcriptomic analysis of DENV2 infection in the presence of 25-HC. Our findings reveal that 25-HC disrupts membrane organization by altering cholesterol distribution, which inhibits the binding of the DENV envelope (E) protein to lipid membranes and prevents subsequent membrane fusion events. Additionally, 25-HC suppresses the expression of key cholesterol biosynthesis genes, disrupting cholesterol homeostasis required for viral replication. Transcriptomic analysis further shows that 25-HC activates genes involved in inflammatory and innate immune responses, including NLRP3 inflammasome and MAPK signaling pathways, enhancing antiviral immunity. Owing to its broad multi-level action, 25-HC exhibits synergistic effects with established broad-spectrum antivirals, underscoring its potential for combination therapies targeting DENV and other enveloped viruses.

## 2 RESULTS

### 2.1 DENV2 inhibition by 25-HC

In order to evaluate the antiviral effects of 25-HC on DENV, we measured the genomic copy number of DENV2 48 hours post-infection, both with and without 25-HC. To minimize 25-HC’s cytotoxic impact, we determined its CC_50_ value (∼180 µM) on Vero E6 cells (Figure S1D) and ensured that we did not exceed 6% of this value. Our experimental findings indicate that an 8-hour pre-treatment of Vero E6 cells with 25-HC prior to infection substantially reduces Dengue viral RNA levels (assessed at 48 hours post-infection, hpi) in a concentration-dependent fashion, yielding up to an 1800-fold reduction at a concentration of 10 µM with an MOI of 0.05 (Figure 1A). A similar inhibitory trend was observed at higher infection titre (0.5 MOI), where treatment with 10 µM 25-HC resulted in ∼50-fold reduction in viral replication. Under similar 25-HC pre-treatment, BHK-21 cells infected with DENV2 (0.05 MOI) exhibited a ∼100-fold reduction. Even for low 25-HC concentration (500 µM), Vero E6, BHK-21 and Huh-7 cells displayed 18-, 8-, and 2-fold, respective reduction in Dengue viral RNA levels (Figure S1C). These observations underscore that 25-HC demonstrates potent antiviral activity against DENV across various cell types, consistent with findings for Zika virus [15]. However, administering 25-HC 8 hours post-infection in Vero E6 cells significantly reduces its potency, with viral RNA levels decreasing less and genomic copies being 10-fold higher compared to pre-treatment at all tested 25-HC concentrations (Figure 1A). This suggests that 25-HC acts in the early stages of viral infection.

**Figure 1:**
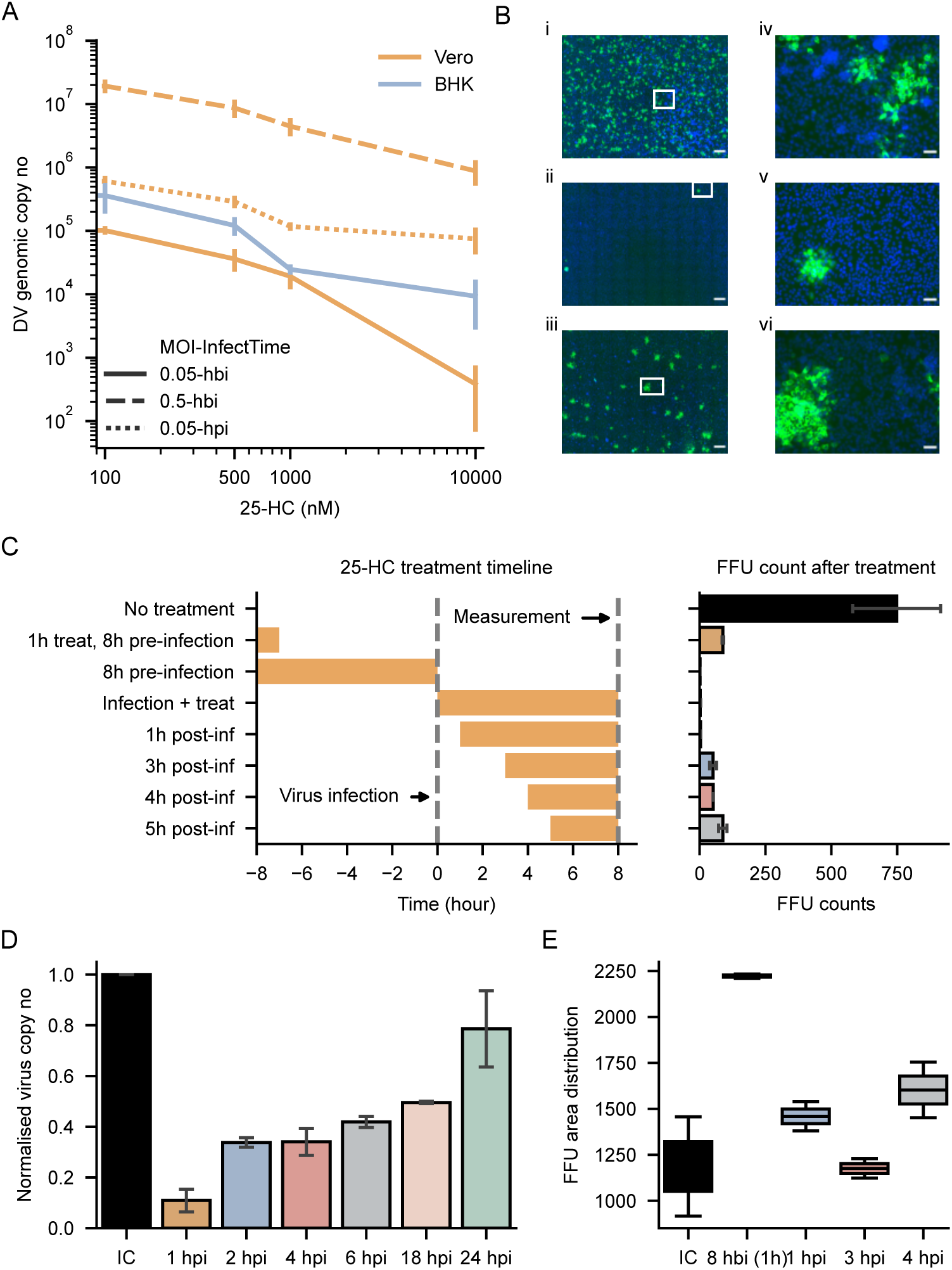
Antiviral activity of 25-HC against Dengue virus 2 (DENV2) (**A**) DENV2 genomic RNA copy numbers (qPCR) measured at 48 hpi upon 25-HC treatment of cells at different concentrations. DENV2 RNA levels for Vero E6 cells upon 25-HC treatment for 8 hours pre-infection or 8 hours post infection are also shown for comparison (n=2, biological replicates). Data are presented as mean ± SEM (B) Immunofluorescence images showing Focus Forming Units (FFU) in DENV-infected Vero E6 cells. (i) Infection control at 0.05 MOI (ii) 8 hrs of pre-treatment with 25-HC before infection (iii) 1 hr pretreatment with 25-HC followed by 7 hr recovery before infection (Scale bars = 500 µm). (iv), (v) and (vi) are cropped zoomed images of the FFUs corresponding to (i), (ii) and (iii), respectively (Scale bars = 50 µm) (**C**) Time-of-addition FFU assay: Left panel shows the schematic representation of time of exposure to 25-HC (1 µM), with respect to infection time of Vero E6 cells with DENV2. FFU counts at 48 hours post-infection (hpi) recorded for each treatment (n=2, biological replicates) are shown in the right panel. Data are presented as mean ± SEM (**D**) DENV genomic copy numbers (qPCR) at 48 hours post infection (hpi) in Vero E6 cells where 25-HC (1 µM) was added at different time periods post infection (**E**) Box plots showing single FFU area distribution (in pixels) for infection control (IC) and with 25-HC treatment for different time intervals corresponding to FFU time-of-addition assay in (**C**) (n=2).

To investigate the mechanism by which 25-HC decreases viral replication, we performed experiments evaluating DENV infection in the presence of 25-HC treatment using time-of-addition FFU assays. A notable decline in virus infectivity was recorded as evidenced by decreased Focus Forming Units (FFUs) when the DENV underwent a pre-incubation with 25-HC (1 µM) for 8 hours (Figure 1B and 1C). Even with just a 1-hour exposure to 25-HC followed by a 7-hour recovery period before DENV infection, a significant reduction in FFU numbers, indicating substantial inhibition, was observed (Figures 1B and 1C). Given that the free 25-HC was rinsed away from the cell culture in this instance, we argue that the cell-membrane-associated 25-HC predominantly mediates the inhibitory effect. However, the application of 25-HC after infection shows a marked reduction in its inhibitory effects, which is consistent with the observations from viral RNA analysis (Figure 1C and 1D). Prolonging the delay in 25-HC addition further reduce its inhibition effectiveness, substantiating its role as an early lifecycle inhibitor (Figure 1C and 1D). In case of late additions of 25-HC, FFU sizes closely resembled those of untreated controls, indicating that viral replication competency remained largely intact and that the principal effect of 25-HC pertains to inhibiting viral entry. However, when cells were pre-treated with 25-HC for a short duration (1 hour) prior to DENV infection resulted in an increase of over 80% in FFU size. This suggests that the signaling induced by 25-HC may interfere with gene expression and cellular metabolism, thereby influencing the replication fitness of the virus in Vero E6 cells (Figure 1B and 1E). These paradoxical observations suggest that 25-HC modulates DENV infection through dual mechanisms: interference with viral entry and early establishment of productive infection while altering cellular phenotype to promote replicative fitness.

### 2.2 25-hydroxycholesterol inhibits Dengue virus membrane fusion

To investigate how 25-HC inhibits DENV, we initially assessed its impact on DENV membrane fusion *in vitro*. We employed fluorescence-based assays to examine virus-liposome membrane fusion, focusing on both lipid and content mixing. Liposomes were prepared with equimolar amounts of POPC, POPG, SM, and CH as controls (quaternary control), and then altered with varying 25-HC concentrations while maintaining total sterol content. Fusion was first measured via dequenching of DiD dye, which fluoresces more when mixed with SUVs of quaternary control composition (POPC:SM:CH:POPG at 1:1:1:1) under pH 5.5 or lower (Figure S2A). The presence of 25-HC in target liposomes resulted in a concentration-dependent reduction in fusion (Figure 2A), with lipid mixing dropping by at least half when over 50% of target cholesterol was substituted with 25-HC, at 12.5% 25-HC (POPC:SM:CH:HC:POPG at 1:1:0.5:0.5:1). To avoid interference in fusion due to DiD in the virus membrane, a FRET pair of NBD-PE (donor) and Rhodamine PE (acceptor) was used to track energy transfer during membrane fusion, showing a similar ∼2-fold decrease when 25-HC was added (Figure 2D). For content mixing, we employed calcein dequenching linked to liposome dye leakage, noting a ∼2-fold decrease in membrane disruption (Figure 2B, S2B, S2C). Further assessment through genome exposure indicated a ∼5-fold decline in content mixing when 12.5% 25-HC was present in target liposome membranes (Figure 2B).

**Figure 2:**
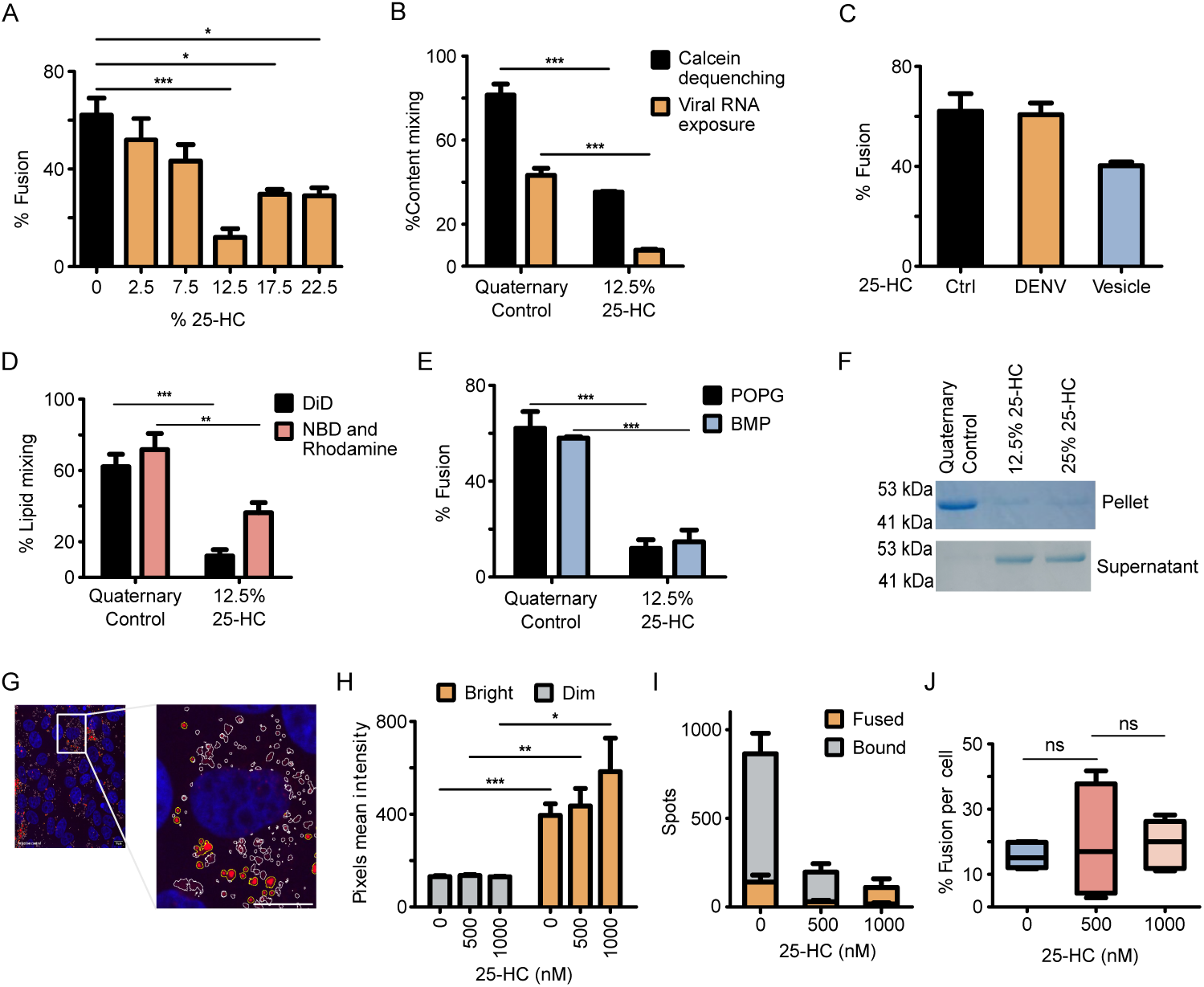
Effect of 25-HC on membrane fusion *in vitro* (**A**) DENV membrane fusion efficiency as a function of percentage of 25-HC of total lipid composition in the vesicle measured from lipid dye dequenching of DiD-labeled virus at pH 5.5. All the quaternary control vesicles (0% 25-HC) contained POPC:POPG:SM:CH lipids in 1:1:1:1 ratio and increasing 25-HC levels are compensated by reducing cholesterol to keep total sterol levels constant (n=3) (**B**) Comparison of content mixing quantified through calcein dequenching and viral RNA genome exposure with or without 12.5% 25-HC in vesicles (n=3) (B) Comparison of membrane fusion efficiency measured by DiD dequenching when either DENV (25-HC: 1 µM for 1 hour) or vesicles (25-HC: 1 µM for 8 hours) are pre-incubated with 25-HC (n=3) (**D**) Comparison of lipid mixing during DENV membrane fusion performed by labeling the target membrane with a FRET pair (NBD and Rhodamine-PE) against DiD dequenching with 12.5% 25-HC in the vesicles (n=3) (**E**) Membrane fusion (DiD dequenching) in the presence of two different anionic lipids (POPG and BMP) with 12.5% 25-HC shows similar reduction in fusion efficiency in both cases (n=3) (**F**) SDS-PAGE analysis of a vesicle sedimentation assay shows reduction in DENV2 envelope protein binding to vesicles with 12.5% and 25% 25-HC at pH 5.5 (**G**) Confocal image showing DiD-labeled DENV (red) binding to Vero E6 cells (nucleus in blue, Hoechst). Zoomed images show assignment of bound (yellow circles) and fused (white circles) viral particles based on intensity analysis (**H**) Mean intensity of bright and dim viral particles quantified from images as shown in **G** panel (N_images_=5) (**I**) Stacked bar chart quantifying total particles per cell under 25-HC pre-treatment (N_images_=5) (**J**) Box plots showing the percentage of fused particles (N_images_=5). The central line marks the median, the box boundaries represent the 25th and 75th percentiles, and whiskers indicate the minimum and maximum values. All data presented as mean ± SEM unless mentioned otherwise. *\*p* < 0.05, *\*\*p* < 0.01, *\*\*\*p* < 0.001, ns: Not Significant; by two-tailed paired t-test (A, B, D, E, and I).

In order to gain a mechanistic understanding of the role of 25-HC in inhibiting fusion, we examined how the envelope protein (E) interacts with model membranes containing varying concentrations of 25-HC, given its established role in virus-target membrane fusion events [6]. Employing a sedimentation assay, we observed that the envelope protein exhibited robust binding with quaternary control liposomes (devoid of 25-HC) at low pH, indicated by a pronounced band in the membrane-bound pellet, while no such band appeared in the unbound supernatant fraction. Conversely, 12.5% and 25% 25-HC markedly reduced E-protein membrane binding, evidenced by weak membrane-associated bands and diminished calcein leakage. (Figures 2F and S2D). The minimal binding phenotype changes observed between 12.5% and 25% 25-HC suggest that 25-HC presence in the model membrane restricts cholesterol availability to the E protein, thereby impeding the fusion process. This finding aligns with recent studies [31] indicating that accessible cholesterol is prevalent in bacterial and viral infections, and 25-HC plays a key role in diminishing the accessible cholesterol pool by mobilizing it away from the host cell membrane surface. Subsequently, to determine if the external uptake of 25-HC might influence membrane fusion, we pre-treated either liposomes or the virus with 1 µM 25-HC and employed the DiD lipid mixing assay to evaluate membrane fusion (Figure 2C). In this scenario, we found approximately 25% reduction in membrane fusion when vesicles were pre-treated with 1 µM 25-HC for 8 hours, whereas no significant reduction was noted when the virus underwent the same pre-treatment. Previous studies have demonstrated that the negatively charged phospholipid bis(monoacylglycerol)phosphate (BMP) is prevalent in late endosomal membranes and is implicated in membrane remodeling and the genesis of intraluminal vesicles in late endosomes [12]. In the present investigation, we formulated liposomes comprising 25% BMPs to analyze membrane fusion, where a 4-fold decrease in membrane fusion was recorded with the presence of 25-HC in the target membrane (Figure 2E), comparable to the effect of POPG in the target membrane.

To examine whether the inhibitory role of 25-HC arises from defect in viral attachment or membrane fusion, we adapted DiD-dequenching virus fusion assay on Vero E6 cell membranes [32]. During DENV’s internalization, membrane fusion occurs between the virus and the host cell within the endosomal compartment, instigated by a pH-dependent alteration in the viral envelope protein [32, 33, 34]. To estimate the level of membrane fusion, we compare the fusion of DiD-labelled virus particles on Vero E6 cell membranes. As a control, we verified that direct labeling with DiD dye of cell membrane does not result in punctate label as observed with DiD labeled viruses (Figure S4). Overall, post receptor binding on the cell plasma membrane, pre-fused viruses display low fluorescence signature per puncta due to quenched DiD fluorescence. However, post viral fusion, dequenching leads to an increased DiD intensity (Figure 2G, 2H and S5A). This enhancement indicates the fusion of viral and endosomal membranes, thereby enabling the estimation of viral fusion events through the ratio of bright to total particle numbers (Figure 2I and S5A, see Methods). We found that pre-treating Vero E6 cells with varying levels of 25-HC for 4 hours did not modify the fusion percentage (Figure 2J), but there was a significant decrease in the total number of DiD-labeled viruses observed on the cell membrane, indicating diminished binding upon 25-HC exposure (Figure 2I). Lack of spread of the DiD dye post fusion (and dequenching) suggests that we are indeed monitoring viral membrane fusion in endosomes (Figure S5B).

Since DENV entry has been shown to be critically dependent on cholesterol and sphingomyelin lipid domains [35], and 25-HC is shown to previously disrupt such microdomains in lipid bilayers [36], we hypothesized that early inhibition of viral entry is driven by 25-HC mediated disruption of cholesterol-rich domains on the cell membrane. This is consistent with observed significant modification of the cholesterol-rich domains upon 25-HC treatment (Figure S3)

### 2.3 Dengue virus suppression in macrophages through 25-hydroxycholesterol induced pathways

Although 25-hydroxycholesterol (25-HC) is well-established as a potent antiviral molecule produced by macrophages (a key cell type infected by DENV) through induction of CH25H[19, 15, 37, 17], the precise mechanisms by which it inhibits DENV remain unclear. To address these gaps, we investigated the impact of 25-HC on DENV infection in macrophages, focusing on its modulation of CH25H expression, innate immune activation, and transcriptional reprogramming, alongside its broader effects on viral replication and host cell pathways. Specifically, 25-HC pre-treatment of RAW 264.7 macrophages for eight hours significantly decreased the intracellular DENV RNA (infection at 0.5 MOI for 24 hrs) by six-fold, mirroring observations in Vero E6 and BHK-21 cells, supporting a cell type-independent antiviral action of 25-HC (Figure 3A). In line with previous studies with other viral pathogens, up-regulation of cholesterol-25-hydroxylase (CH25H) was prominently observed (Figure 3B) after DENV infection in various cell lines, including Vero E6 (Figure 3E) and BHK-21 cells (Figures S6A). Notably, this response was particularly pronounced, with an approximate 80-fold increase in RAW 264.7 cells as further confirmed by gel electrophoresis of qPCR products and corresponding western blot analysis (Figure 3B and Figures S6B,S6C). This suggests that CH25H upregulation during flavivirus infection[15], modulates its own expression and activates antiviral pathways across diverse cell types. Consistent with lower DENV levels, 25-HC pretreatment of macrophages resulted in lowering CH25H expression during DENV infection, possibly due to reduced viral replication associated activation of ISG pathways (Figure 3B). It is likely that macrophages can be primed to a broad antiviral state due to the activation ISG pathways during infections. To test this, non-viral activation of RAW 264.7 cells was performed using lipopolysaccharide (LPS for 4 hours) prior to DENV infection. This approach was sufficient to prime the macrophages to an antiviral state leading to a stark reduction in the DENV growth (Figure 3C). This enhanced antiviral state of the cell was confirmed by the elevated levels of CH25H expression following LPS exposure, underscoring the role of a broader innate immune response in antiviral protection (Figure 3D). We hypothesized that exogeneous addition of 25-HC might also activate components of this broad ISG primed antiviral cell state and play a role in DENV inhibition. Indeed, 25-HC alone was able to induce CH25H expression in macrophages and Vero E6 cells, implying a self-reinforcing activation loop that could prime an antiviral state of the cell (Figures 3B,3E).

**Figure 3:**
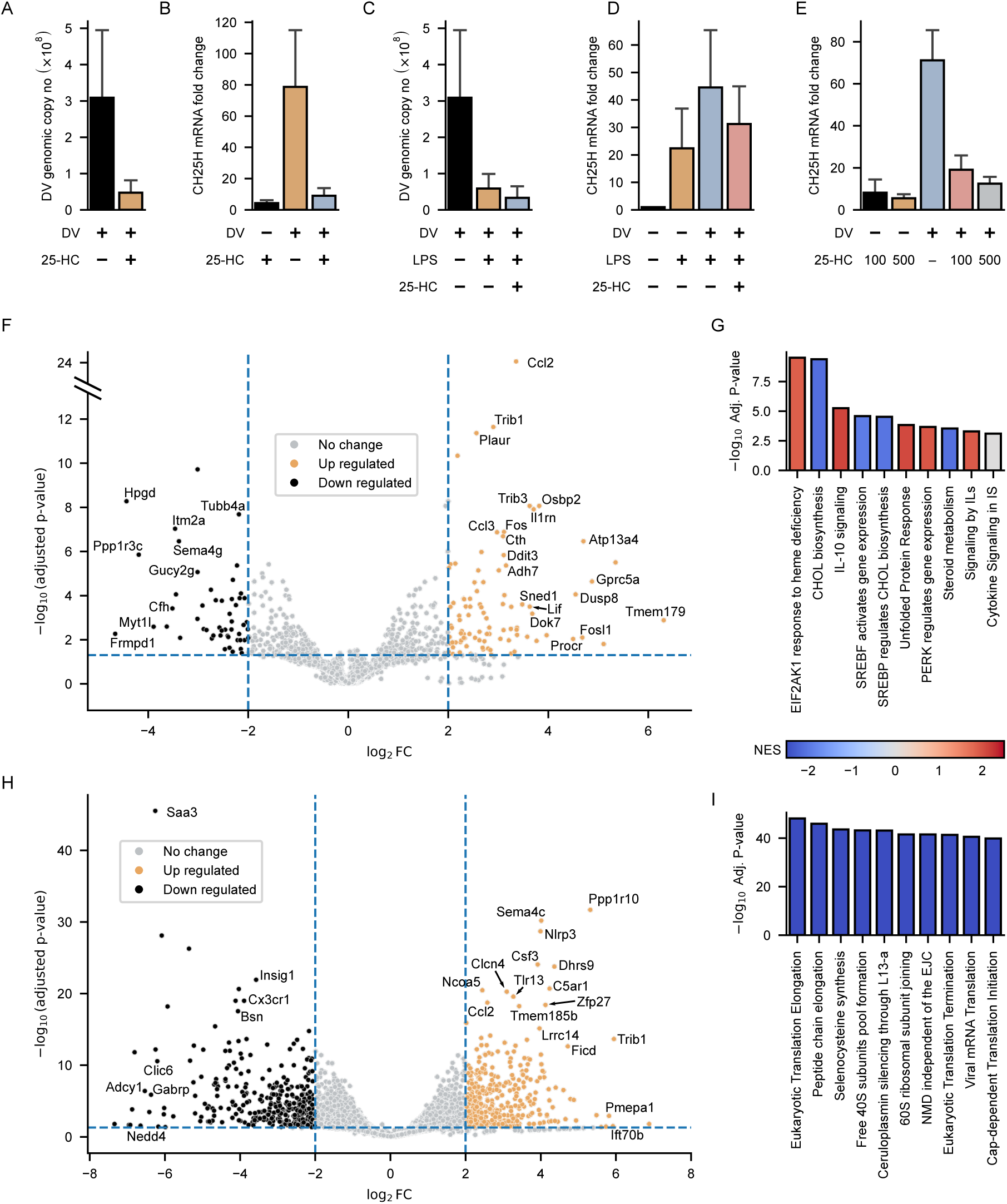
Differential gene expression in RAW 264.7 cell during DENV infection and 25-HC exposure (**A**) DENV genomic copy number in RAW 264.7 cells following pre-incubation with 25-HC (1 µM, 24 hrs). (**B**) CH25H gene expression normalized to β-actin in response to LPS treatment, with or without DENV infection or 25-HC treatment, in RAW 264.7 cells. (**C**) CH25H expression in RAW 264.7 cells in response to DV infection, 25-HC treatment, and LPS pre-treatment. (**D**) DENV genomic copy number in RAW 264.7 cells following LPS pre-treatment. (**E**) CH25H gene expression normalized to β-actin in response to DENV infection under varying concentrations of 25-HC in Vero E6 cells. All data presented above is mean ± SEM (n=3) (**F**) Volcano plot showing the differentially regulated genes and (**G**) Reactome enrichment analysis in RAW 264.7 cells only treated with 25-HC with respect to control (n=2). Similarly, (**H**) and (**I**) show respective plots for RAW 264.7 cells pre-treated with 25-HC before infection against no pre-treatment before infection (n=2). Color represents Normalised Enrichment Score (NES) from GSEA analysis.

To explore the regulatory effects of 25-HC on DENV (DENV) through host cell gene expression, we conducted RNA-Seq analysis to assess variations in host gene expression levels following DENV infection, both with and without the presence of 25-HC. In alignment with previous findings [38], DENV infection in innate immune cells initiates pathways connected to pro-inflammatory response, immune activation, chemotaxis, and stress response (Figure S7A,S7C). Pre-exposure to 25-HC alters DENV infection with notable changes in gene expression, particularly in pathways involving adaptive immune responses, inflammatory response, and innate immune activation (Figure S7B,S7D). Specifically, a significant upregulation of crucial immune and inflammatory response genes such as NLRP3, CCL2, CCL7, and CSF3 is observed in cells infected with DENV and pre-treated with 25-HC in comparison to cells with DENV infection alone (Figure 3H). To determine if this reaction stems solely from the viral infection or if 25-HC enhances the innate immune response to the infection, we analyzed gene expression differences between DENV-infected cells with and without 25-HC (Figure 3H). The amplified transcriptional response aligned with findings from gene ontology (GO) enrichment analysis, where immune and inflammatory activities are significantly enriched, highlighting the potent immune activation caused by 25-HC during DENV infection (Figure S7B).

As illustrated in Figure 3F, treatment with 25-HC induced a pronounced shift in the transcriptomic profile towards a pro-inflammatory and stress-responsive state. This shift was characterized by the upregulation of key MAP Kinase pathway genes (e.g., TRIB1, FOS), core stress-response regulators (DDIT3, OSBP2), chemokines (CCL2, CCL3), and cytokines belonging to the IL-6 family (OSM, LIF), collectively indicating coordinated activation of MAPK and chemokine signaling pathways. Reactome enrichment analysis of significantly upregulated genes (NES > 1, FDR < 0.05) confirmed robust activation of pathways related to ”Cytokine Signaling in Immune System” and ”Signaling by Interleukins,” along with pathways involved in endoplasmic reticulum stress (”PERK Regulated Gene Expression”) and lipid metabolism regulation mediated by SREBF (Figure 3G).

DENV infection triggers a robust response in macrophages, characterized by major changes in innate immune signaling, lipid metabolism, and other critical pathways that contribute to viral replication and disease pathogenesis (Figure S8B). Transcriptomic studies reveal that DENV infection upregulates interferon-stimulated genes (ISGs) such as IFITM1, MX2, and OAS1A, while simultaneously suppressing key antiviral pathways, including STAT1 phosphorylation, to evade immune responses. Notably, key interferon-stimulated genes such as MX2, RSAD2, and IRF7 were significantly upregulated, indicating activation of the type I interferon signaling axis. Infected macrophages produce elevated levels of pro-inflammatory cytokines like TNF-α, IL-36, and IL-1β, fueling systemic inflammation (Figure S7C). This upregulation of inflammatory responses has also been attributed to vascular leakage in severe dengue cases [39]. In parallel, the induction of CCL22 points to enhanced chemokine-mediated immune signaling. DENV infection also induces significant metabolic reprogramming in macrophages, particularly in lipid and cholesterol pathways [40]. DENV relies on cholesterol-enriched membranes inside cell to assemble its replication complexes and upregulates key cholesterol biosynthesis genes such as HMGCR and LDLR to meet this requirement, with lipid droplets providing a potential reservoir for membrane lipids. [41, 42]. CRISPR knockout studies have further identified host factors critical for viral replication, including SREBF2, a regulator of cholesterol biosynthesis, and ER-associated proteins like SEC61A1. Beyond metabolism, DENV modulates autophagy, oxidative stress, and apoptotic pathways, with increased expression of genes like ATG5 and NQO1 and dysregulation of apoptotic regulators such as BAX and BCL2.

Comparative analysis of dengue-infected RAW 264.7 cells, with or without prior 25-HC treatment, revealed that 25-HC pre-treatment significantly altered the transcriptional landscape upon dengue infection. Notably, there was enhanced expression of innate immune sensors and effectors, alongside MAPK regulators, suggesting that 25-HC pre-treatment potentiates TLR- and inflammasome-driven innate immune responses, cytokine–chemokine signaling, and MAPK pathways in dengue-infected macrophages (Figure 3H). 25-HC pre-treatment substantially downregulated pathways associated with protein synthesis, affecting stages of host and viral protein translation, including cap-dependent initiation, 40S subunit joining, peptide-chain elongation, and translation termination whereas there was enhancement in virus infection and pathways associated with translation in case of DENV infection (Figure 3I). Reactome pathway analysis revealed significant enrichment of viral infection pathways, infectious disease, and multiple facets of protein synthesis in DENV–infected macrophages. These included key stages of host and viral translation such as eukaryotic translation initiation, cap-dependent initiation, and formation of the 40S ribosomal subunit pool (Figure S8B). However, pre-treatment with 25-HC (DENV+HC condition) markedly suppressed these translation-associated pathways (Figure S8C). Specifically, we observed broad downregulation of pathways involved in cap-dependent initiation, 60S subunit joining, peptide-chain elongation, and translation termination, indicating that 25-HC disrupts both host and viral protein synthesis machinery. These findings suggest that while DENV infection enhances the host’s translational landscape to facilitate viral replication, 25-HC counteracts this by targeting key stages of the protein synthesis process.

### 2.4 Modulation of lipid and cholesterol homeostasis by 25-HC during Dengue infection

One of key transcriptional perturbations induced by 25-HC was dysregulation of lipid metabolism genes particularly those involved in cholesterol biosynthesis (Figure 3G). To validate 25-HC induced lipid reorganization in macrophages, we imaged and observed dramatic upregulation of lipid droplets in macrophages post 25-HC exposure (Figures 4B, 4C and 4E). Further, when we imaged labelled cholesterol (TF-Chol) distribution, we note a prominent reorganization and co-localization of cholesterol with lipid droplets (Figures 4C, 4D and 4H) in contrast to untreated cells (Figure 4A). Conversely, cells incubated with labelled hydroxycholesterol (TF-25-HC, green) and stained for lipid droplets (red) exhibited significantly less overlap between the signals (Figures 4E, 4F and 4H).

**Figure 4:**
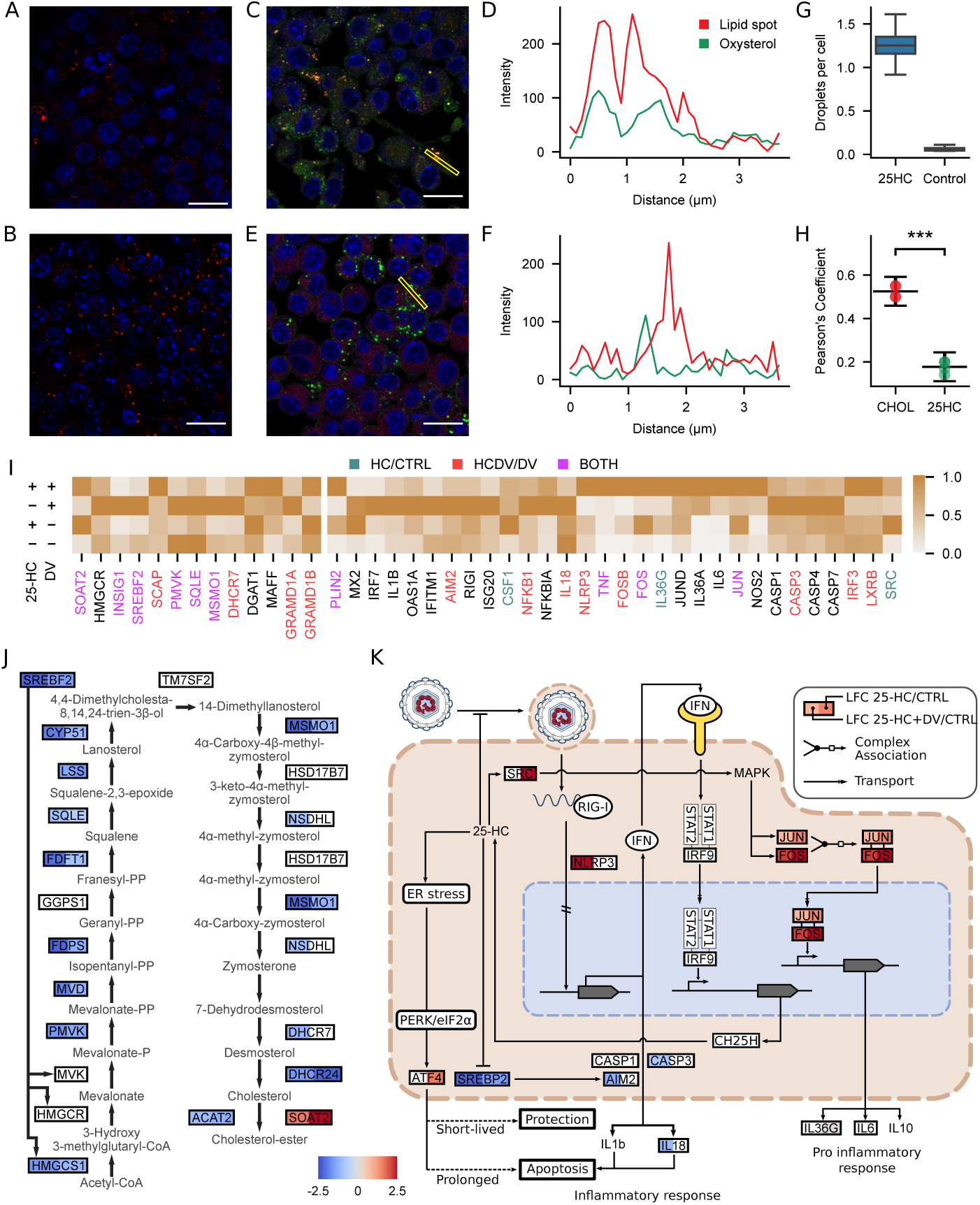
25-HC modulates cholesterol metabolism and host antiviral immune signalling in DENV-infected macrophages. Confocal microscopy images of (**A**) Untreated RAW 264.7 cells stained with LipidSpot dye to visualize lipid droplets (red). (**B**) RAW 264.7 cells treated with unlabelled 25-HC (1 µM, 4 h) and stained for lipid droplets (red). (**C**) RAW 264.7 cells incubated with unlabelled 25-HC (1 µM, 4 h) lipid droplets (red), and fluorescently labelled cholesterol (green). (**D**) Line-scan intensity profile from the region marked in (**C**), showing partial overlap of cholesterol with lipid droplets. (**E**) RAW 264.7 cells incubated with fluorescently labelled 25-HC (1 µM, 4 h; green) and stained for lipid droplets (red). Scale bars: 8 µm.(**F**) Line-scan analysis of the region marked in (**E**), showing lack of overlap between labelled 25-HC and lipid droplets. (**G**) Quantification of lipid droplet number per cell in control vs 25-HC–treated RAW 264.7 cells. High threshold for Lipid Spot dye was used to avoid counting background noise as LDs. Error bars represent min and max data points (n=4) (**H**) Pearson’s correlation coefficient for TopFluor cholesterol vs lipid droplet colocalization and TopFluor 25-HC vs lipid droplets. Data presented as mean ± SEM (n=4) (**I**) Heatmap of differentially expressed genes across untreated RAW 264.7, DENV infected (DV), 25-HC treated and DENV infection post 25-HC treatment. Select cholesterol biosynthesis genes, interferon-stimulated genes (ISGs), and immune response genes are shown for brevity. Expression values are scaled per gene, with significant changes in test/control comparison indicated by colored gene labels (n=2) (**J**) Schematic representation of the cholesterol biosynthesis pathway showing metabolic nodes altered by DENV infection and/or 25-HC treatment (n=2) (**K**) Integrated schematic illustrating the effect of 25-HC on antiviral immune signaling pathways during DENV infection, including modulation of MAPK signaling, ER stress response, and proinflammatory cytokine production. Colormap indicates the log2FoldChange in RNA levels NOTE: log_2_ FoldChange > 2 condition was droppped in making this figure. *\*\*\*p* < 0.001 by paired t-test.

Transcriptomic analysis provides clear insights into the significant alterations observed in lipid metabolism following 25-hydroxycholesterol (25-HC) treatment (Figure 4I and 4J). Specifically, we observed robust downregulation of genes central to cholesterol biosynthesis, including the master regulatory transcription factor SREBF2, along with its downstream target enzymes SQLE, PMVK, MSMO1, and DHCR7 upon 25-HC exposure. Conversely, the expression of INSIG1 was markedly increased during DENV infection, likely serving as a negative feedback regulator that binds to SCAP, thus preventing excessive cholesterol synthesis. Notably, increased SCAP expression specifically in the presence of both 25-HC and DENV infection further suggests an intrinsic feedback mechanism aimed at tightly regulating cholesterol synthesis via modulation of the SREBP pathway. Given that DENV infection typically elevates SREBF2-driven cholesterol synthesis, the presence of 25-HC during infection effectively counteracts this virus-induced upregulation (Figure S9). The results suggest that while 25-HC suppresses cholesterol synthesis, it enhances cholesterol esterification through SOAT2 upregulation. This suppression is wide-spread across the entire mevalonate to cholesterol pathway (Figure 4J) suggesting that direct 25-HC interaction with the SREBP2 might be responsible for this effect [27, 25]. Similarly, genes associated with lipid droplet formation, notably PLIN2 DGAT1, GRAMD1B, SOAT2 [43, 44, 45, 46] were markedly upregulated upon 25-HC exposure confirming enhanced lipid droplet biogenesis observed by imaging (Figure 4B). This effect can lead to a possible increase in DENV replication and pathogenesis within host cells, suggesting that 25-HC may also act as a minor agonist for DENV growth. Consistent with transcriptomic reprogramming reported by [47], we found 25-HC treatment can induce robust ER stress by activating the integrated stress response pathway, which includes upregulation of key stress response genes and selective translation of stress-related transcripts via EIF2AK1 (HRI)-mediated response to heme deficiency, unfolded protein response, and PERK-regulated gene expression pathways, ultimately reprogramming macrophage gene expression toward an ER stress phenotype (Figure 3G). We observed selective upregulation of GRAMD1B in 25-HC–treated and 25-HC + DENV-infected macrophages, likely reflecting IL-6–driven activation of GRAMD1B to restore lipid homeostasis via JAK/STAT signaling [48] (Figure 4I), whereas GRAMD1A induction in DENV-only infection aligns with its reported role as an immunecheckpoint–associated biomarker, suggesting virus-driven reprogramming of lipid-immune regulatory axes [49].

Apart from repressing SREBP-driven sterol synthesis, our RNA-seq analysis revealed that 25-hydroxycholesterol (25-HC) treatment significantly upregulated the nuclear receptor Liver X Receptor beta (LXRβ) and the cholesterol-transport protein Oxysterol-binding protein (OSBP). Previous studies have demonstrated that LXR activation facilitates cholesterol efflux by inducing genes involved in cholesterol transport and esterification [50]. Furthermore, OSBP is known to mediate non-vesicular cholesterol trafficking between cellular membranes, contributing to the efficient redistribution and esterification of intracellular cholesterol [51]. Therefore, our findings suggest that 25-HC not only suppresses cholesterol biosynthesis pathways but also activates LXR signaling and OSBP-mediated cholesterol trafficking, collectively promoting enhanced cholesterol esterification and lipid-droplet formation in macrophages.

### 2.5 Priming of macrophage antiviral pathways by 25-HC

DENV infection is known to profoundly reprogram macrophage function, driving changes in innate immune signaling, lipid metabolism, and other critical pathways that contribute to viral replication and disease pathogenesis. Previously, transcriptomic studies have revealed that DENV infection upregulates interferon-stimulated genes (ISGs) such as IFIT1, MX1, and OAS1, while simultaneously suppressing key antiviral pathways, including STAT1 phosphorylation, to evade immune responses [52].

In contrast, 25-HC treatment established an alternative antiviral state, independent of IFN-I induction. Specifically, 25-HC upregulated CSF1 and transcription factors Fos/Jun, both of which are tightly linked to MAPK–ERK1/2 signalling in macrophages. LXRs, known sensors of oxysterols, may also contribute to this regulation by cross-talking with MAPK cascades, thereby amplifying AP-1 (Fos/Jun) activity.

When combined with DENV infection, 25-HC further potentiated innate immune effector pathways, as reflected in the induction of NLRP3, TNF-α, NOS2, IL-36α, and IL-36γ, suggesting a switch toward a heightened proinflammatory and antiviral macrophage state. Thus, while DENV primarily exploits and manipulates IFN signalling, 25-HC primes macrophages through CSF1–MAPK–AP-1–driven programs that compensate for viral immune evasion and mount an alternative antiviral defense. 25-HC has been documented to amplify inflammatory reactions in influenza virus infections by facilitating AP-1 recruitment and activating ER stress mechanisms [30]. Our research, however, elucidates a more intricate function of 25-HC in modulating immune responses triggered by DENV in murine macrophages. The expression patterns of Interferon Stimulated Genes (ISGs), along with chemokines and cytokines observed here, highlight the significant immuno-regulatory effects of 25-HC during DENV infection. Notably, ISGs such as ISG15, ISG20, IFITM family, and Viperin are strongly induced by DENV, yet their expression is notably diminished following 25-HC pre-treatment, indicating a specific modulation or suppression of interferon signalling. Intriguingly, 25-HC pre-treatment shows involvement with components of the AP-1 transcription factor family, including Fos-b, Jun, and Jund, which selectively activate AP-1 dimers, thus enhancing the expression of key inflammatory mediators like IL36a, IL6, IL36g, TNF, and certain CCL chemokines (CCL2, CCL3, CCL4). Moreover, 25-HC pretreatment significantly elevates NLRP3 expression, while pro-inflammatory and apoptotic factors such as AIM2, Caspase 1, and Caspase 3 are upregulated during DENV infection, resulting in increased cytokine production like IL1b and IL18 (Figure 4I). Yet, the upregulation of CSF or anti-inflammatory cytokine IL10 during 25-HC pretreatment signifies its role in the modulation of interferon signalling and inflammasome activation, emphasizing a complex immune regulatory function of 25-HC during viral infections. Figure 4K presents a network model depicting how 25-HC can redirect immune responses from excessive inflammation towards a more balanced antiviral defence, potentially enhancing both innate and adaptive immune systems.

### 2.6 Synergistic action of 25-HC with direct-acting and host-acting antivirals

While 25-HC displayed potent antiviral activity, induction of ER stress and associated cytotoxicity at higher concentrations can limit its utility in clinical settings. However, its broad and multi-stage inhibitory effect also makes it amenable to synergistic combination with other antivirals that target distinct steps in the virus life cycle [53]. Thus, we next explored combinatorial strategies for 25-HC with different class of antivirals to determine whether synergy could reduce the effective dose of these drugs while enhancing antiviral efficacy [54]. To elucidate the molecular pathway targeted by 25-HC, we opted for drugs with proven anti-flaviviral effects to understand that drug interactions can either amplify or diminish their overall efficacy. Overall, we examined the synergistic effects of 25-HC in combination with antivirals that inhibit membrane fusion (Picolinic acid[55]), disrupt endosomal acidic environments (Chloroquine [56]), or interfere with viral replication (Remdesivir[57] and K22 [58] (N-3- [4-(4-bromophenyl)-4-hydroxypiperidin-1-yl]-3-oxo-1-phenylprop-1-en-2-yl benzamide)). The assessment of these interactions was executed using a 3×3 dose matrix format, testing combinations of 25-HC with the other drugs (see Methods).

Prior research has demonstrated that Remdesivir is capable of inhibiting RNA-dependent RNA polymerases (RdRps) across various RNA virus families, including Filoviridae, Coronaviridae, and Flaviviridae [57]. Our findings indicated that the combination of 25-HC and Remdesivir (RD) significantly decreased viral titers, achieving reductions up to 50% greater than either compound alone (Figure 5A). This is clearly evident from Synergy Index (SnI) values that ranges from 0.33-0.56 at 500 nM 25-HC (Figure 5B). We posit that the dual targeting of distinct stages in the viral lifecycle—entry (25-HC) and replication (RD) —amplifies the antiviral efficacy in combination. Furthermore, 25-HC primes the host immune system by activating innate immune pathways, including the NLRP3 inflammasome and MAPK signaling, which enhance the antiviral state of the host cells. This immune modulation likely augments the efficacy of direct-acting antivirals, creating a robust antiviral environment. Notably, the synergy observed allows for a reduction in the effective doses of both 25-HC and the companion drugs, mitigating cytotoxicity and improving the therapeutic index.

**Figure 5:**
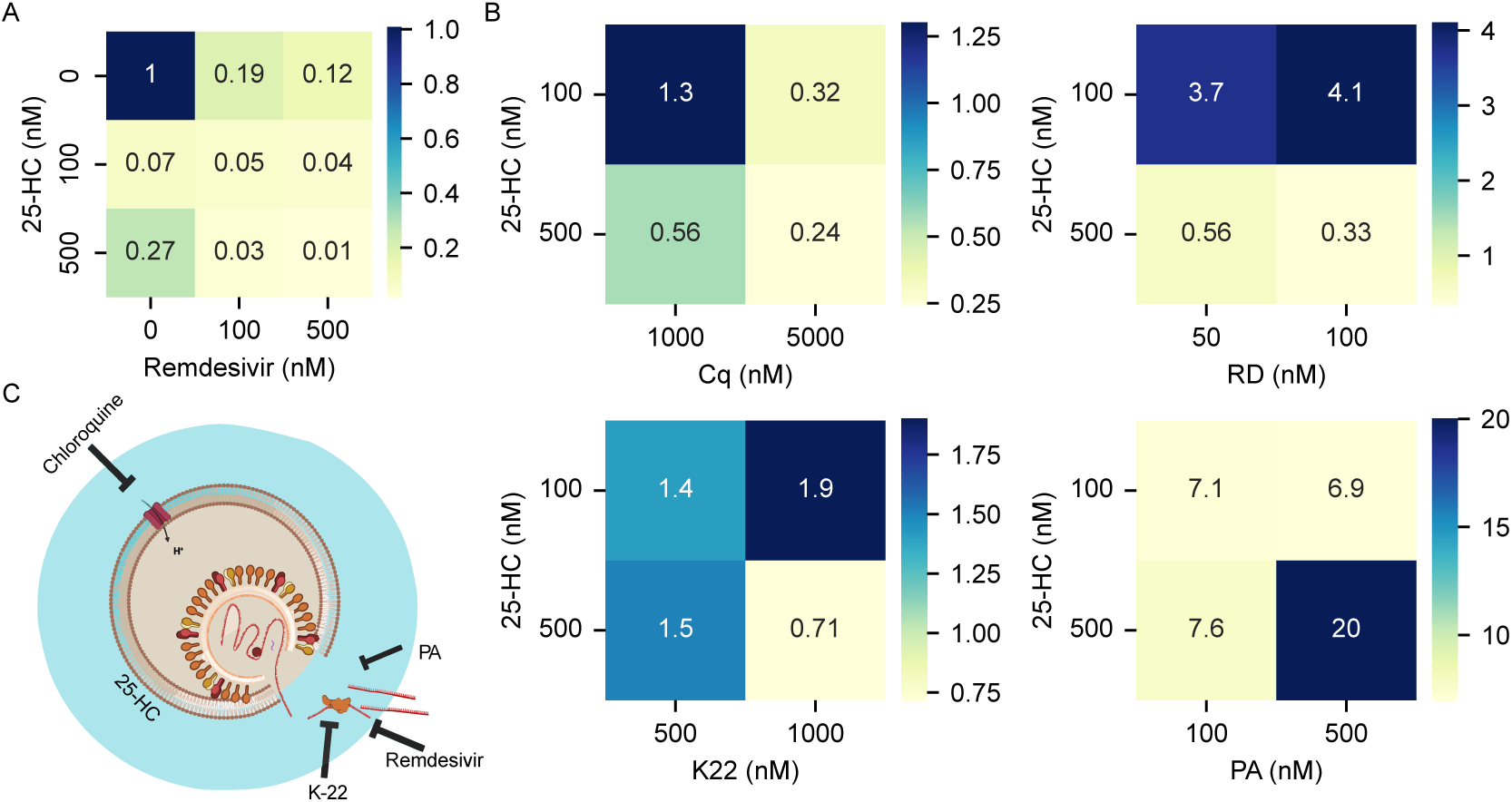
Drug synergy between 25-HC and other antivirals including Chloroquine (Cq), K-22, Remdesivir (RD) and Picolinic acid (PA) (**A**) Virus copy number normalised to untreated control for the combination of Remdesivir and 25-HC (**B**) Comparative synergy between 25-HC and Chloroquine, Remdesivir, K-22, and Picolinic acid for Vero E6 cells for different drug concentrations is shown as a heatmap (n=3) (**C**) Schematic representation illustrating how each drug inhibits virus infection by interfering at different stages of the viral lifecycle.

On the hand, K-22, that inhibits several Flaviviridae members by disrupting the formation of membranebound replication compartments [58], displayed positive synergy (SnI < 1) only at the highest concentrations tested (Figure 5B). Additionally, we explored the synergistic effects of host-targeting drugs, including Chloroquine, an established amine acidotropic agent. This compound has been demonstrated to inhibit various RNA viruses, like SARS-CoV-2, by obstructing the terminal glycosylation of cellular receptors [59], and it impacts Zika and DENV by increasing endosomal pH [56]. The synergistic use of 25-HC and chloroquine at higher drug concentrations resulted in a striking 80% reduction in viral load compared to their individual effects (Figure 5B and S10C) leading to strong postive synergy across most combinations tested (SnI ranging from 0.24-1.3).

Conversely, Picolinic acid, known for its wide-ranging antiviral effects against enveloped viruses like SARS-CoV-2, Influenza A virus, and flaviviruses, acts by interfering with viral fusion to cellular membranes and impeding endocytosis [55], appears to be an ineffective companion in combination therapy with 25-HC. The pairing of 25-HC and Picolinic acid resulted in dose-dependent viral suppression without significant augmented reduction when combined (Figure S10A and 5B). In fact, 25-HC displayed poor synergistic efficacy in presence of PA at all concentrations tested (SnI = 6.9-20). The data indicates that having a strong overlap in mechanistic actions does not markedly enhance synergy, possibly due to both drugs converging on the pathway aimed at inhibiting membrane fusion (Figure 5C). Figure 5C illustrates that the mechanism of drug interactions highlights the molecular foundation underlying the observed synergistic antiviral effects.

## 3 DISCUSSION

Although 25-HC is recognised for its wide-ranging suppression of infections caused by enveloped and non-enveloped viruses, the detailed molecular pathways underlying its extensive antiviral efficacy are not fully elucidated. Our findings reveal that 25-HC exerts its antiviral effects against DENV through a dual mechanism: (1) biophysical disruption of membrane architecture and cholesterol distribution that manifests itself in reduced viral membrane fusion, and (2) selective regulation of immune signaling pathways to mitigate viral pathogenesis and inflammation. Given its extensive and occasionally opposing effects on cellular physiology, 25-HC also triggers stress-related pathways and lipid droplet accumulation, which might undermine its efficacy as an antiviral. Consequently, we propose that 25-HC efficacy is most effectively utilised when incorporated into combination therapy with other antiviral drugs.

The biophysical disruption of membrane architecture by 25-HC is a key mechanism underlying its antiviral activity. Cholesterol-rich lipid rafts are essential for DENV entry, as they facilitate the binding of the viral envelope (E) protein to host cell membranes and subsequent viral fusion [60]. Incorporation of 25-HC into target membranes significantly alters cholesterol distribution and availability, impairing lipid raft organization [61]. This disruption reduces the binding affinity of the DENV E protein, thereby inhibiting viral fusion and entry. Notably, our cellular analyses indicated that although 25-HC disrupts interactions occurring at the membrane level, it does not interfere with the process of endocytic membrane fusion after the virus has adhered to the surface of the host cell membrane. Moreover, the disruption of viral membrane fusion resulting from 25-HC incorporation within target membranes, irrespective of receptor interaction (demonstrated by *in vitro* fusion experiments), indicates that changes in membrane structure and composition are crucial for its antiviral activity. This inhibitory effect is more pronounced when 25-HC is administered prior to DENV infection, highlighting its prophylactic potential. These findings suggest that 25-HC primarily targets the early stages of viral entry by modulating membrane composition, rather than interfering with endocytic processes.

In addition to cholesterol’s role in viral entry, DENV infection enhances cholesterol synthesis, which is crucial for its replication and assembly processes [62]. Our observations indicate that DENV infection leads to upregulation of key cholesterol biosynthesis genes, including SREBF2, HMGCR, and SQLE, reflecting increased cholesterol production. This enhancement likely provides cholesterol-rich membranes required during the viral life cycle. Concurrently, the upregulation of Insig1 indicates a host regulatory mechanism in response to the excessive cholesterol accumulation which is potentially harmful to the host cell. On the other hand, 25-HC significantly inhibits cholesterol synthesis by diminishing the expression of HMGCR, SREBF2, and SQLE during DENV infection. We also observe the differential regulation of SREBP1 and SREBP2 due to exposure to 25-HC. While 25-HC reduces SREBP2 activation via INSIG1, thus limiting cholesterol biosynthesis and its availability for viral replication, SREBP1 remains relatively unaffected owing to its lesser sensitivity to sterols and regulation by INSIG2. This imbalance leads to an increase in fatty acid and triglyceride synthesis, resulting in lipid droplet accumulation which can serve as a platform for viral replication. Interestingly, SCAP, a critical regulator of SREBP activation, is upregulated in presence of 25-HC, possibly reflecting a feedback mechanism to cope with low levels of SREBP2 and INSIG1. This upregulation of SCAP selectively favours SREBP1 activation, enhancing lipid droplet formation even as cholesterol synthesis is restricted. Thus this metabolic alteration exerts a dual outcome: restricting cholesterol-dependent viral activities while simultaneously providing triglyceride-rich resources in the form of lipid droplets that are conducive to viral replication. Therefore, potential therapeutic strategies focusing on SCAP or SREBP1 could be exploited to boost the antiviral efficacy of 25-HC.

Beyond its direct antiviral properties, 25-HC influences the host’s immune modulation. DENV infection enhances ISG expression through the JAK-STAT signaling cascade, including the overexpression of CH25H gene in macrophages. However, pre-exposure to 25-HC reduces this activation, thereby potentially mitigating excessive inflammatory response. Instead, 25-HC enhances AP-1 transcriptional activity, promoting a modulated immune response that can limit viral pathogenesis and inflammation. It seems that 25-hydroxycholesterol (25-HC) establishes a distinct, alternate antiviral state against DENV infection that differs significantly from the host’s natural immune response. While natural DENV infection promotes a broad, interferon-centric response marked by the strong upregulation of interferon-stimulated genes (ISGs) like ISG15, IFITM family, and Viperin, our data from Figure 3F and 3H reveals that 25-HC treatment effectively dampens the expression of these same ISGs. Instead, 25-HC redirects the autocrine feedback loop to induce a more targeted inflammatory and antiviral state, evidenced by the significant upregulation of specific genes such as NLRP3, CCL2, CCL7, and CSF3. This suggests that 25-HC’s therapeutic effect is not simply an amplification of the natural immune response, but a unique modulation that establishes an environment less conducive to viral replication by suppressing conventional ISG pathways and, in conjunction with its effects on TLR-mediated signaling, promotes a distinct inflammatory profile.

The antiviral potential of 25-HC is limited by poor pharmacokinetics, off-target effects like cytotoxicity and inflammation, and challenges in systemic delivery due to its hydrophobic nature. Additionally, its long-term impact on lipid homeostasis, host stress pathways and immune regulation remains unclear, limiting its widespread usage. However, we demonstrate that combination therapy with drugs acting on direct viral targets displayed positive synergy with 25-HC. For example, the synergy between 25-HC and Remdesivir likely arises from their complementary actions where 25-HC disrupts viral replication complexes by cholesterol dysregulation and interferon response, while, Remdesivir directly inhibits the viral RdRP, reducing the viral burden and enhancing the overall effectiveness of the immune response. Conversely, Piconolic acid displays significant negative drug interaction with 25-HC, consistent with their shared role in membrane fusion inhibition. This observed synergy with early and late-life cycle stage inhibitors as well as between IFN and direct-acting antivirals is consistent with predictions from viral dynamics mathematical models[53, 63]. Therefore, strategic combinations of antivirals with innate immune modulators, such as 25-HC, represent a promising new avenue for the development of potent and well-tolerated antiviral therapies.

## 4 METHODS

### 4.1 Cell culture

BHK-21 (ATCC CRL-1587), Vero E6 (ATCC CRL-1587), RAW 264.7 (a generous gift from Dr. Siddharth Jhunjhunwala), Huh-7 (a generous gift from Dr. Soumitra Das) cells were maintained in Dulbecco’s Modified Eagle Medium (DMEM, Gibco Thermo-Fisher Scientific), supplemented with 10% fetal bovine serum (Thermo-Fisher Scientific), 15 mM HEPES, 200 mM glutamine, 100 mM non-essential amino acids, and penicillin-streptomycin 100 U/ml at 37*^◦^*C, 5% CO_2_.

### 4.2 Virus production

The plasmid pFK-DV encoding a synthetic copy of full-length genomic sequence DENV2 strain 16681 was used to generate the DENV as previously reported [64]. Briefly, viral RNA was synthesized from the linearized plasmid by in vitro transcription (HiScribeTM SP6 RNA Synthesis Kit) followed by capping. RNA was purified by Sodium-acetate precipitation and transfected to the BHK-21 cells by lipofectamine Messenger-Max (Thermo-Fisher Scientific) and the supernatant containing infectious virus was harvested and cleared from the cell debris by low-speed centrifugation. Passage 1 (P1) virus was produced by infecting Vero E6 cells with the supernatant collected above. The virus particles were collected and purified from cell debris by low-speed centrifugation. This supernatant containing P1 virus was further used for infecting cells in the presence or absence of different concentrations of 25-HC.

### 4.3 Focus Forming Unit (FFU) assay

Approximately 0.05×10^6^ Vero E6 cells were seeded in 24 well plates to grow and reach a confluency of 70%. Cells were then infected with serial threefold dilutions of DENV and incubated for 1 hour at 37*^◦^*C to allow adsorption. The inoculum was then removed and the cells were overlaid with 1.5% carboxymethylcellulose (CMC) in DMEM supplemented with 2% FBS. After 48 of incubation, cells were fixed with 4% para formaldehyde and permeabilized with 0.1% Triton X-100. Infected foci were detected using a virus-specific primary antibody 4G2 with 1:500 dilution (Catalog No. GTX57154) and secondary antibody of Goat anti-Mouse IgG (H+L) Cross-Adsorbed Secondary Antibody, Alexa Fluor™ 488 (A-11001) with the concentration of 1 µg/mL. The infected cell clusters (focus) were captured with an inverted microscope (Evos M7000 microscope). Briefly, 100 tiled images were captured at 20X magnification and merging them into a single merged image (∼ 0.09 cm^2^) (Figure S1A). Fluorescent images from the fluorescence detection channel (488 nm excitation/500-550 nm emission) channel, indicative of viral infection, was used to quantify the number of foci-forming units (Figure S1B). Images were processed using code that implemented a intensity threshold combined with a minimum area size threshold to identify and select objects, followed by the creation of a mask. Refinement steps included dilation, erosion, opening, and closing operations, with all parameters standardized and consistently applied across all images. The code is available at https://github.com/Debayani12/25HC.

### 4.4 Quantification of Dengue virus genomic RNA and CH25H gene expression with qPCR

To quantify DENV genomic RNA from cell culture supernatants, viral particles were first concentrated using polyethylene glycol 8000 precipitation, followed by resuspension in Tricine buffer (20 mM Tricine, pH 7.8, 140 mM NaCl, and 0.005% Pluronic F-127). Total RNA was extracted using RNAiso Plus reagent (Takara, 9108/9109) according to the manufacturer’s instructions. Complementary DNA (cDNA) synthesis was performed using SuperScript™ IV Reverse Transcriptase (Thermo Fisher Scientific, 18090010) with random hexamer primers. For quantitative PCR, 100 ng of cDNA was amplified using Phusion High-Fidelity DNA Polymerase (NEB, M0530S), 10 µM each of forward (5′GTCCCCTTCTCAGGCAGAATGAGCC-3′) and reverse (5′-GTTCGTGTCTCCAGTCCCATTCCC-3′) primers, and EvaGreen dye in a 1× reaction buffer. Viral genome copy number was calculated using a standard curve generated from serial dilutions of a plasmid containing the full-length DENV2 genomic sequence.

Quantitative PCR was performed to measure CH25H gene expression relative to *β*-actin using TB Green (Takara). For RAW 264.7 cells, the primers used were forward 5′-CCATAAGGAC-AAGGGAGGCG-3′ and reverse 5′-GTCCGGGTGGATCTTGTAGC-3′; for Vero E6 cells, forward 5′-CTGACCTTCCACGT-GGTCAA-3′ and reverse 5′-GTGTGTGAAGTACGGAGCGA-3′; and for BHK-21 cells, forward 5′-ACCCGAATTGCTCACTCCTC-3′ and reverse 5′-CCTCGCTACAGAATCGTGCT-3′.

### 4.5 Time of Addition assay

For the Time of Addition FFU assay, approximately 0.05×10^6^ Vero E6 cells were seeded in 24-well plates and cultured to ∼70% confluency. Post infection in presence of different time periods with 25-HC, cells were fixed, stained and imaged as per FFU procedure described above. Cells were either pre-treated with 25-HC for varying durations or infected with DENV at a multiplicity of infection (MOI) of 0.05, followed by post-infection treatment with 25-HC (as per the FFU assay protocol described above). For genomic RNA quantification, ∼ 0.3×10^6^ Vero E6 cells were seeded in 6-well plates and grown to ∼70% confluency. These cells were treated with 25-HC either prior to or following infection with DENV at MOIs of 0.05 or 0.5. Viral genome copy number was determined by qPCR as described above.

### 4.6 Confocal imaging of RAW 264.7 cells with lipid dyes and/or CTXB and colocalization analysis

RAW 264.7 macrophages were cultured to approximately ∼60-70% confluency and treated with 1 µM 25-hydroxycholesterol (25-HC) for 4 hours. For lipid treatment and colocalization analysis, cells were treated with 25-HC either in combination with TopFluor Cholesterol (Avanti, 810255) or TopFluor 25-HC (Avanti, 810299) (1 µM). After washing with PBS, cells were stained with LipidSpot™ (Biotium, 70069-T) dye for 30 minutes to visualize neutral lipids for colocalization analysis. For confocal imaging, cells were also incubated with fluorescently labeled cholera toxin B subunit (CTXB; 1 µg/mL, 30 minutes at 37 °C, Invitrogen). Confocal images for both experiments were acquired on a Leica SP8 microscope using a 63× oil-immersion objective.

### 4.7 Image analysis

Lipid droplets were quantified from confocal images using code available at https://github.com/Debayani12/25HC. The code was run with intensity thresholding (5xStdev above background) and size filters to segment and count droplets per cell.

For CTXB labeled RAW cell images, texture analysis was applied to pre-and post-treatment micrographs to quantify treatment-induced differences without requiring precise segmentation. We used Local Binary Patterns (LBP) [65] and Gray-Level Co-occurrence Matrices (GLCM) [66], as these descriptors have been shown to be effective for histopathology image classification [67]. LBP encodes local micro-patterns like edges, spots, corners; shifts in its histogram indicate changes in fine-scale roughness and pattern diversity and are robust to monotonic intensity variations. GLCM characterizes pairwise gray-level relationships at specified distances and orientations, yielding interpretable metrics, e.g., contrast as a proxy for roughness/edge density, where higher value of contrast represents more edges/texture/roughness. Together, LBP captures local texture while GLCM summarizes second-order, direction- and scale-sensitive structure, revealing both fine- and mesoscale changes with respect to granularity, clustering, and alignment between the two image.

### 4.8 Transcriptomic analysis

RAW 264.7 cells were seeded at a density of 0.1×10^6^ cells in 6-well plates prior to the day of experiment and treated with 1 µM 25-HC for eight hours, followed by wash and DENV infection for 24 hours. After treatment, cells were washed with PBS and lysed with TRIzol reagent (Invitrogen). mRNA libraries were sequenced using Illumina NovaSeq X plus (MedGenome) to generate 2×150 bp reads/sample. The raw RNA-seq reads were trimmed to remove adapter sequences and low-quality bases (Phred score < 20) using Trim Galore![68]. Quality assessment of the trimmed reads was performed using FastQC[69]. The raw reads were aligned to GRCm39 mouse genome [70] with STAR aligner [71] with default parameters. The counts of aligned reads per gene were determined using featureCounts in paired-end mode, focusing on the gene meta feature level [72], with multimapping reads being excluded. After counting, PyDESeq2 (a Python implementation of DESeq2) was used for differential gene expression analysis [73]. Genes with adjusted p-value of less than 0.05 and absolute log_2_ FoldChange > 2 (after LFC shrinkage) were considered as differentially expressed genes. These genes were used as input for statistical overrepresentation test for gene ontology with PantherDB [74, 75, 76]. Background genes for GO enrichment analysis were used from an reference RNAseq dataset [77]. The FDR correction method was employed to adjust for multiple comparisons. Gene Set Enrichment Analysis was conducted using the GSEApy [78] package, utilizing data from Gene Ontology [76]. Reactome pathway information was acquired from reactome.org [79], and a pathway over-representation analysis was executed locally with GSEApy, considering all genes participating in a pathway as the background.

### 4.9 Synergy assays

About 0.05 × 10^6^ Vero E6 cells and 0.04 × 10^6^ BHK-21 cells were cultured in a 24-well plate to achieve approximately 70% confluency. We performed checkerboard assays wherein at least two concentrations of antiviral compounds such as Chloroquine (Sigma-C6628-25G), Picolinic acid (Sigma-P42800), remdesivir (MedChem-HY-104077) and K-22 (N-3-[4-(4-bromophenyl)-4-hydroxypiperidin-1-yl]-3-oxo-1-phenylprop-1-en-2-ylbenzamide) (Chemdiv 4295-0370) were tested in combination with two concentrations of 25-HC in a 3×3 matrix format of concentrations below their respective IC_50_ values for 4 hours. Post drug treatment cells were infected with DENV at 0.05 MOI. After 48 hours of infection, culture supernatants were collected, and viral RNA levels were quantified using qPCR.

We determined the synergy score using the equation S = Y_c_/(Y_1_ × Y_2_), where Y_c_ represents the relative reduction in viral RNA when using the drug combination, and Y_1_ and Y_2_ correspond to the reductions seen with each drug individually. A score of S < 1 was interpreted as evidence of positive synergy.

### 4.10 Cell viability (MTT) assay

0.01×10^6^ Vero E6 cells were seeded into 96-well plates and allowed to adhere overnight. The following day, cells were treated with increasing concentrations of 25-hydroxycholesterol (25-HC), and cytotoxicity was evaluated using the MTT assay (M6494, Thermo Fisher Scientific) according to the manufacturer’s instructions.

### 4.11 Western Bolt analysis and Envelope protein vesicle binding assay

RAW 264.7 cells were cultured to ∼70% confluency, treated for 48 hrs with 1 µM 25-HC, and subjected to western blotting using anti-CH25H and anti-*β*-actin antibodies (Invitrogen).

For the Envelope protein binding assay, multilamellar vesicles (MLVs) were prepared as described previously [80]. Lipid–protein binding analysis was performed by incubating 300 µL of 1 mM MLVs with DENV envelope protein (final protein concentration of 5 µM) at 37*^◦^*C for 30 minutes with gentle mixing. The vesicle–protein suspension was then centrifuged at 22,000 × g for 30 minutes to separate bound and unbound proteins. The supernatant was carefully collected, and the pellet was resuspended in 300 µL phosphate-buffered saline (PBS). Both fractions (supernatant and pellet) were analyzed by SDS-PAGE to determine protein distribution. Protein bands were visualized using Coomassie Brilliant Blue staining to identify the fraction enriched with the DENV envelope protein.

### 4.12 Preparation of labelled virus for fluorescence dequenching assay

The virus supernatant was collected 72 hours after the infection, and the virions were pelleted by ultracentrifugation at 4 *^◦^*C for 15 hours for 30000x*g*. The virus pellet was resuspended in a buffer containing 20 mM Tricine (N- (2-hydroxy-1,1 bis (hydroxymethyl) ethyl) glycine) pH 7.8, 140 mM NaCl and 0.005% Pluronic F-127 (Tricine buffer) and 500 nM DiD as previously described. The virus was further purified over an Opti-prep density gradient (32000 rpm at 4*^◦^*C for 3 hours) with 15%, 25%, 40% and 60% density levels. The fraction ranging from 25% to 40% was harvested, aliquoted, and preserved at –80*^◦^*C. The virus was separated and purified from Opti-prep using a 100 kDa filter (Amicon ultra 100 kDa centrifugal filter unit). The viruses labeled with DiD were stored at 4*^◦^*C and utilized for the dequenching assay within 2 days.

### 4.13 Labelled virus cell fusion assay

0.01×10^6^ Vero E6 cells were seeded in µ-Slide 8 Well coverslip (ibidi 80826) and either treated with different concentrations of 25-HC for four hours or without pre-treatment, washed, kept on ice for 10 minutes and infected with 5 MOI of DiD labelled virus for one 1.5 h at 37 *^◦^*C. The virus inoculum was then removed, fixed with 4% paraformaldehyde (PFA) and incubated with DAPI, washed and finally mounted on glass slide. Images were acquired using Andor Dragonfly Spinning Disk confocal microscope. The virus quantification in relation to the DAPI-positive cells was carried out using ImageJ/Fiji software.

### 4.14 Labeled vesicle preparation for fusion assays

Small Uni-lamellar Vesicles (SUV) were prepared with the lipid composition of POPC:SM:CH: POPG as 1:1:1:1 or of POPC:SM:CH: HC: POPG at a ratio of 1:1:0.5:0.5:1. Briefly, lipids were mixed in chloroform that was made to evaporate under nitrogen to form a thin film in the glass vial. Residual chloroform was made to evaporate by keeping the samples in vacuum overnight. The lipid films were then hydrated in PBS, or fluorescent dye calcein (80 mM) in phosphate buffer saline (PBS) or SYTO RNA Select (10 µM). Liposomes were also prepared in varying concentrations of 25-HC, altering cholesterol. Liposomes were prepared at pH 5 and 7.5 respectively.

### 4.15 Content mixing assays

Liposomes encapsulating calcein were prepared according to a previously established protocol [80]. The intensity was measured in a kinetic manner using excitation/emission wavelengths of 495 and 515 nm with a 10 nm bandwidth, beginning from the baseline intensity (I_0_). Subsequently, 5 μl of virus (diluted 10-fold from stock) was added, and the contents were mixed to measure the intensity (I_t_). Once the signal reached saturation, 1% TritonX was introduced to fully dequench the dye, achieving the maximal intensity (I_max_). The degree of liposome content leakage (and membrane fusion) was evaluated using the dequenching ratio, by comparing the dequenching due to viral fusion with that elicited by Triton treatment, as

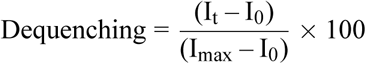

RNA genome exposure during virus membrane fusion with vesicles is monitored with STYO RNA Select loaded vesicles. Fluorescence was measured of the vesicles before (A_0_), and after (A_i_), fusion with excitation/emission maxima set at ∼490/530 nm (10 nm window). To normalize the level of fluorescence change upon RNA binding, a total of 1 μg of IVT RNA was incubated with the SYTO RNA Select dye and dye fluorescence before (R_0_), and after (R_i_) was measured. The viral RNA fluorescence change (considered content mixing efficiency) during virus membrane fusion was normalized by comparing the signal change to the fluorescence change observed upon binding of the SYTO RNA Select dye to naked IVT RNA.

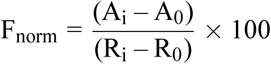

### 4.16 Lipid mixing assays

Fusion was monitored via (i) DiD dequenching and (ii) FRET-based assays using liposomes labeled with NBD/Rhodamine-PE. The virus liposome fusion was monitored at pH 5.5 and pH 7.5 respectively. A liposome fusion reaction was performed on a fluorescence spectrophotometer with excitation and emission wavelengths as 467 nm and 590 nm. 95 μl of the liposome was added into the cuvette (F_0_) followed by the addition of virus (F_1_) and 1% TritonX respectively. Full fusion was measured as a function of the decrease of the acceptor fluorescence (Rhodamine-PE) signal due to fusion. Signals were normalized to the lowest fluorescence produced by the detergent solubilization and the fusion curves were plotted as a percentage of fusion efficiency considering maximum fusion efficiency with 1% TritonX 100 as 100%(F_max_).

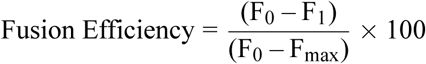

All bulk fluorescence measurements were conducted on Cary Eclipse Fluorescence Spectrometer (Agilent).

### 4.17 Data and code availability

RNA-seq data, both raw and processed, produced as part of this research, along with the supplementary analysis code for RNA-seq processing and ontology analysis, can be obtained by contacting Rahul Roy at. The bespoke code utilized for image analysis and synergy computation is accessible at https://github.com/Debayani12/25HC.

## Acknowledgements

We thank Dr Ralf Bartenschlager (University of Heidelberg) for generously providing the pFK-DV plasmid, used to generate DENV. We thank Dr Saumitra Das for providing Huh-7 cell line and Dr Siddharth Jhunjhunwala for providing Raw 264.7 cell line. This work was supported by funding from the Human Frontier Science Program (HFSP), the National Postdoctoral Fellowship (NPDF), and the Ministry of Human Resource Development (MHRD), Government of India. We gratefully acknowledge MedGenome for assistance with mRNA sequencing. We also thank our laboratory members for their helpful discussions and technical support during the study. We acknowledge the Department of Chemical Engineering for infrastructural support and IISc Bio-imaging facility for confocal microscopy.

## 4.18 Author Contributions

RR, A.S., D.C. did conceptualization of the study, methodology, formal analysis, writing of the original draft. D.C., A.S, M.K., D.T., A.P. conducted experiments and/or performed analysis. R.R, M.K, D.C, A.S., D(B)C analyzed data, provided key technical and conceptual advice. D.C, A.S., M.K, A.P., D.T., R.R. provided comments and suggestions on the final manuscript.

## 4.19 Competing Interests

The authors declare no conflict of interest.

## Notes

### Competing Interest Statement

The authors have declared no competing interest.

### Summary of Updates

This version of the manuscript includes details of lipid droplet generation by 25-HC, membrane cholesterol distribution inferring how membrane accessible cholesterol accumulate in lipid droplets.

## References

[1] Julia Noack and Shaeri Mukherjee. ”make way”: Pathogen exploitation of membrane traffic. Current Opinion in Cell Biology, 65:78–85, Mar 2020. doi: 10.1016/j.ceb.2020.02.011.

[2] Judith M. White, Amanda E. Ward, Laura Odongo, and Lukas K. Tamm. Viral membrane fusion: A dance between proteins and lipids. Annual Review of Virology, 10(1):139–161, Sep 2023. doi: 10.1146/annurev-virology-111821-093413.

[3] Rakesh Kulkarni, Erik A. C. Wiemer, and Wen Chang. Role of lipid rafts in pathogen-host interaction - a mini review. Frontiers in Immunology, 12:815020, Jan 2022. doi: 10.3389/fimmu.2021.815020.

[4] Margaret Kielian and Félix A. Rey. Virus membrane-fusion proteins: more than one way to make a hairpin. Nature Reviews Microbiology, 4(1):67–76, Jan 2006. doi: 10.1038/nrmicro1326.

[5] Stephen C Harrison. Viral membrane fusion. Nature Structural & Molecular Biology, 15(7):690–698, July 2008. ISSN 1545-9993, 1545-9985. doi: 10.1038/nsmb.1456.

[6] Yorgo Modis, Steven Ogata, David Clements, and Stephen C. Harrison. Structure of the dengue virus envelope protein after membrane fusion. Nature, 427(6972):313–319, January 2004. ISSN 0028-0836, 1476-4687. doi: 10.1038/nature02165. URL https://www.nature.com/articles/nature02165.

[7] A. C. Walls, Y. J. Park, M. A. Tortorici, A. Wall, A. T. McGuire, and D. Veesler. Structure, function, and antigenicity of the sars-cov-2 spike glycoprotein. Cell, 181(2):281–292.e6, Apr 2020. doi: 10.1016/j.cell.2020.02.058.

[8] Jung Shim, Jinhee Kim, Tanel Tenson, Ji-Young Min, and Denis Kainov. Influenza Virus Infection, Interferon Response, Viral Counter-Response, and Apoptosis. Viruses, 9(8):223, August 2017. ISSN 1999-4915. doi: 10.3390/v9080223. URL https://www.mdpi.com/1999-4915/9/8/223.

[9] V. A., et al. Kostyuchenko. Structure of the thermally stable zika virus. Nature, 533:425–428, Apr 2016. doi: 10.1038/nature17994.

[10] Stephen C. Harrison. Viral membrane fusion. Virology, 479-480:498–507, may 2015. ISSN 00426822. doi: 10.1016/j.virol.2015.03.043. URL https://linkinghub.elsevier.com/retrieve/pii/S004268221500183X.

[11] Jason Mercer, Mario Schelhaas, and Ari Helenius. Virus Entry by Endocytosis. Annual Review of Biochemistry, 79(1):803–833, June 2010. ISSN 0066-4154, 1545-4509. doi: 10.1146/annurev-biochem-060208-104626. URL https://www.annualreviews.org/doi/10.1146/annurev-biochem-060208-104626.

[12] Elena Zaitseva, Sung-Tae Yang, Kamran Melikov, Sergei Pourmal, and Leonid V. Chernomordik. Dengue Virus Ensures Its Fusion in Late Endosomes Using Compartment-Specific Lipids. PLoS Pathogens, 6(10):e1001131, October 2010. ISSN 1553-7374. doi: 10.1371/journal.ppat.1001131. URL https://dx.plos.org/10.1371/journal.ppat.1001131.

[13] Kai Simons and Elina Ikonen. Functional rafts in cell membranes. Nature, 387(6633):569–572, June 1997. ISSN 0028-0836, 1476-4687. doi: 10.1038/42408. URL https://www.nature.com/articles/42408.

[14] Nicholas S. Heaton and Glenn Randall. Multifaceted roles for lipids in viral infection. Trends in Microbiology, 19(7):368–375, July 2011. ISSN 0966842X. doi: 10.1016/j.tim.2011.03.007. URL https://linkinghub.elsevier.com/retrieve/pii/S0966842X11000576.

[15] Chunfeng Li, Yong-Qiang Deng, Shuo Wang, Feng Ma, Roghiyh Aliyari, Xing-Yao Huang, Na-Na Zhang, Momoko Watanabe, Hao-Long Dong, Ping Liu, Xiao-Feng Li, Qing Ye, Min Tian, Shuai Hong, Junwan Fan, Hui Zhao, Lili Li, Neda Vishlaghi, Jessie E. Buth, Connie Au, Ying Liu, Ning Lu, Peishuang Du, F. Xiao-Feng Qin, Bo Zhang, Danyang Gong, Xinghong Dai, Ren Sun, Bennett G. Novitch, Zhiheng Xu, Cheng-Feng Qin, and Genhong Cheng. 25-Hydroxycholesterol Protects Host against Zika Virus Infection and Its Associated Microcephaly in a Mouse Model. Immunity, 46(3):446–456, March 2017. ISSN 1097-4180. doi: 10.1016/j.immuni.2017.02.012.

[16] David Lembo, Valeria Cagno, Andrea Civra, and Giuseppe Poli. Oxysterols: An emerging class of broad spectrum antiviral effectors. Molecular Aspects of Medicine, 49:23–30, June 2016. ISSN 1872-9452. doi: 10.1016/j.mam.2016.04.003.

[17] S. Y., et al. Liu. Interferon-inducible cholesterol-25-hydroxylase broadly inhibits viral entry by production of 25-hydroxycholesterol. Immunity, 38(1):92–105, Jan 2013. doi: 10.1016/j.immuni.2012.11.005.

[18] B. Gomes, S. Gonçalves, A. Disalvo, A. Hollmann, and N. C. Santos. Effect of 25-hydroxycholesterol in viral membrane fusion: Insights on hiv inhibition. Biochimica et Biophysica Acta (BBA) - Biomembranes, 1860(5):1171–1178, May 2018. doi: 10.1016/j.bbamem.2018.02.001.

[19] Ruochen Zang, James Brett Case, Eylan Yutuc, Xiucui Ma, Sheng Shen, Maria Florencia Gomez Castro, Zhuoming Liu, Qiru Zeng, Haiyan Zhao, Juhee Son, Paul W. Rothlauf, Alex J. B. Kreutzberger, Gaopeng Hou, Hu Zhang, Sayantan Bose, Xin Wang, Michael D. Vahey, Kartik Mani, William J. Griffiths, Tom Kirchhausen, Daved H. Fremont, Haitao Guo, Abhinav Diwan, Yuqin Wang, Michael S. Diamond, Sean P. J. Whelan, and Siyuan Ding. Cholesterol 25-hydroxylase suppresses SARS-CoV-2 replication by blocking membrane fusion. Proceedings of the National Academy of Sciences, 117(50):32105–32113, December 2020. ISSN 0027-8424, 1091-6490. doi: 10.1073/pnas.2012197117. URL https://pnas.org/doi/full/10.1073/pnas.2012197117.

[20] Yongzhi Chen, Shanshan Wang, Zhaohong Yi, Huabin Tian, Roghiyh Aliyari, Yanhua Li, Gang Chen, Ping Liu, Jin Zhong, Xinwen Chen, Peishuang Du, Lishan Su, F. Xiao-Feng Qin, Hongyu Deng, and Genhong Cheng. Interferon-Inducible Cholesterol-25-Hydroxylase Inhibits Hepatitis C Virus Replication via Distinct Mechanisms. Scientific Reports, 4(1):7242, December 2014. ISSN 2045-2322. doi: 10.1038/srep07242. URL https://www.nature.com/articles/srep07242.

[21] Valeria Cagno, Andrea Civra, Daniela Rossin, Simone Calfapietra, Claudio Caccia, Valerio Leoni, Nicholas Dorma, Fiorella Biasi, Giuseppe Poli, and David Lembo. Inhibition of herpes simplex-1 virus replication by 25-hydroxycholesterol and 27-hydroxycholesterol. Redox Biology, 12: 522–527, August 2017. ISSN 22132317. doi: 10.1016/j.redox.2017.03.016. URL https://linkinghub.elsevier.com/retrieve/pii/S2213231717301659.

[22] A., et al. Civra. 25-hydroxycholesterol and 27-hydroxycholesterol inhibit human rotavirus infection by sequestering viral particles into late endosomes. Redox Biology, 19:318–330, Oct 2018. doi: 10.1016/j.redox.2018.09.003.

[23] David R. Bauman, Andrew D. Bitmansour, Jeffrey G. McDonald, Bonne M. Thompson, Guosheng Liang, and David W. Russell. 25-Hydroxycholesterol secreted by macrophages in response to Toll-like receptor activation suppresses immunoglobulin A production. Proceedings of the National Academy of Sciences of the United States of America, 106(39):16764–16769, September 2009. ISSN 1091-6490. doi: 10.1073/pnas.0909142106.

[24] M., et al. Blanc. The transcription factor stat-1 couples macrophage synthesis of 25-hydroxycholesterol to the interferon antiviral response. Immunity, 38(1):106–118, Jan 2013. doi: 10.1016/j.immuni.2012.11.004.

[25] A., et al. Reboldi. 25-hydroxycholesterol suppresses interleukin-1-driven inflammation downstream of type i interferon. Science, 345(6197):679–684, Aug 2014. doi: 10.1126/science.1254790.

[26] Ying Liu, Zhuo Wei, Ye Zhang, Xingzhe Ma, Yuanli Chen, Miao Yu, Chuanrui Ma, Xiaoju Li, Youjia Cao, Jian Liu, Jihong Han, Xiaoxiao Yang, and Yajun Duan. Activation of liver X receptor plays a central role in antiviral actions of 25-hydroxycholesterol. Journal of Lipid Research, 59 (12):2287–2296, December 2018. ISSN 00222275. doi: 10.1194/jlr.M084558. URL https://linkinghub.elsevier.com/retrieve/pii/S0022227520341547.

[27] J. D. Horton, J. L. Goldstein, and M. S. Brown. Srebps: activators of the complete program of cholesterol and fatty acid synthesis in the liver. Journal of Clinical Investigation, 109(9):1125– 1131, May 2002. doi: 10.1172/jci200215593.

[28] H., et al. Merta. Spatial proteomics of er tubules reveals clmn, an er-actin tether at focal adhesions that promotes cell migration. Cell Reports, 44(4):115502, Apr 2025. doi: 10.1016/j.celrep.2025.115502.

[29] X. Y. Dong, S. Q. Tang, and J. D. Chen. Dual functions of insig proteins in cholesterol homeostasis. Lipids in Health and Disease, 11:173, Dec 2012. doi: 10.1186/1476-511X-11-173.

[30] E. S., et al. Gold. 25-hydroxycholesterol acts as an amplifier of inflammatory signaling. PNAS, 111 (29):10666–10671, Jul 2014. doi: 10.1073/pnas.1404271111.

[31] Michael E. Abrams, Kristen A. Johnson, Sofya S. Perelman, Li shu Zhang, Shreya Endapally, Katrina B. Mar, Bonne M. Thompson, Jeffrey G. McDonald, John W. Schoggins, Arun Radhakrishnan, and Neal M. Alto. Oxysterols provide innate immunity to bacterial infection by mobilizing cell surface accessible cholesterol. Nature Microbiology, 5:929–942, 7 2020. ISSN 20585276. doi: 10.1038/s41564-020-0701-5.

[32] Hilde M. Van Der Schaar, Michael J. Rust, Barry-Lee Waarts, Heidi Van Der Ende-Metselaar, Richard J. Kuhn, Jan Wilschut, Xiaowei Zhuang, and Jolanda M. Smit. Characterization of the Early Events in Dengue Virus Cell Entry by Biochemical Assays and Single-Virus Tracking. Journal of Virology, 81(21):12019–12028, November 2007. ISSN 0022-538X, 1098-5514. doi: 10.1128/JVI.00300-07. URL https://journals.asm.org/doi/10.1128/JVI.00300-07.

[33] Luke H Chao, Jaebong Jang, Adam Johnson, Anthony Nguyen, Nathanael S Gray, Priscilla L Yang, and Stephen C Harrison. How small-molecule inhibitors of dengue-virus infection interfere with viral membrane fusion. eLife, 7:e36461, July 2018. ISSN 2050-084X. doi: 10.7554/eLife.36461. URL https://elifesciences.org/articles/36461.

[34] Luke H Chao, Daryl E Klein, Aaron G Schmidt, Jennifer M Peña, and Stephen C Harrison. Sequential conformational rearrangements in flavivirus membrane fusion. eLife, 3:e04389, December 2014. ISSN 2050-084X. doi: 10.7554/eLife.04389. URL 10.7554/eLife.04389.

[35] Juan Fidel Osuna-Ramos, José Manuel Reyes-Ruiz, and Rosa Maria Del Ángel. The Role of Host Cholesterol During Flavivirus Infection. Frontiers in Cellular and Infection Microbiology, 8:388, November 2018. ISSN 2235-2988. doi: 10.3389/fcimb.2018.00388. URL https://www.frontiersin.org/article/10.3389/fcimb.2018.00388/full.

[36] Marco M. Domingues, Bárbara Gomes, Axel Hollmann, and Nuno C. Santos. 25-Hydroxycholesterol Effect on Membrane Structure and Mechanical Properties. International Journal of Molecular Sciences, 22(5):2574, March 2021. ISSN 1422-0067. doi: 10.3390/ijms22052574. URL https://www.mdpi.com/1422-0067/22/5/2574.

[37] Mathieu Blanc, Wei Yuan Hsieh, Kevin A Robertson, Kai A Kropp, Thorsten Forster, Guanghou Shui, Paul Lacaze, Steven Watterson, Samantha J Griffiths, Nathanael J Spann, Anna Meljon, Simon Talbot, Kathiresan Krishnan, Douglas F Covey, Markus R Wenk, Marie Craigon, Zsolts Ruzsics, Jürgen Haas, Ana Angulo, William J Griffiths, Christopher K Glass, Yuqin Wang, and Peter Ghazal. The transcription factor STAT-1 couples macrophage synthesis of 25-hydroxycholesterol to the interferon antiviral response. Immunity, 38(1):106–118, January 2013. doi: 10.1016/j.immuni.2012.11.004.

[38] Caroline Fernandes-Santos and Elzinandes Leal De Azeredo. Innate Immune Response to Dengue Virus: Toll-like Receptors and Antiviral Response. Viruses, 14(5):992, May 2022. ISSN 1999-4915. doi: 10.3390/v14050992. URL https://www.mdpi.com/1999-4915/14/5/992.

[39] Gathsaurie Neelika Malavige and Graham S Ogg. Pathogenesis of vascular leak in dengue virus infection. Immunology, 151(3):261–269, jul 2017.

[40] Joao Palma Pombo and Sumana Sanyal. Perturbation of intracellular cholesterol and fatty acid homeostasis during flavivirus infections. Front. Immunol., 9, June 2018.

[41] Shizhan Ma, Wenxiu Sun, Ling Gao, and Shudong Liu. Therapeutic targets of hypercholesterolemia: HMGCR and LDLR. Diabetes Metab. Syndr. Obes., 12:1543–1553, August 2019. doi: 10.2147/DMSO.S219013.

[42] Rushika Perera, Catherine Riley, Giorgis Isaac, Amber S Hopf-Jannasch, Ronald J Moore, Karl W Weitz, Ljiljana Pasa-Tolic, Thomas O Metz, Jiri Adamec, and Richard J Kuhn. Dengue virus infection perturbs lipid homeostasis in infected mosquito cells. PLoS Pathog., 8(3):e1002584, March 2012. doi: 10.1371/journal.ppat.1002584.

[43] Avery L McIntosh, Subramanian Senthivinayagam, Kenneth C Moon, Shipra Gupta, Joel S Lwande, Cameron C Murphy, Stephen M Storey, and Barbara P Atshaves. Direct interaction of plin2 with lipids on the surface of lipid droplets: a live cell FRET analysis. Am. J. Physiol. Cell Physiol., 303(7):C728–42, October 2012. doi: 10.1152/ajpcell.00448.2011.

[44] Lívia Teixeira, Filipe S Pereira-Dutra, Patrícia A Reis, Tamires Cunha-Fernandes, Marcos Y Yoshinaga, Luciana Souza-Moreira, Ellen K Souza, Ester A Barreto, Thiago P Silva, Hugo Espinheira- Silva, Tathiany Igreja, Maísa M Antunes, Ana Cristina S Bombaça, Cassiano F Gonçalves-de Albuquerque, Gustavo B Menezes, Eugênio D Hottz, Rubem F S Menna-Barreto, Clarissa M Maya-Monteiro, Fernando A Bozza, Sayuri Miyamoto, Rossana C N Melo, and Patrícia T Bozza. Prevention of lipid droplet accumulation by DGAT1 inhibition ameliorates sepsis-induced liver injury and inflammation. JHEP Rep., 6(2):100984, February 2024. doi: 10.1016/j.jhepr.2023.100984.

[45] Diana Acosta Ingram, Emir Turkes, Tae Yeon Kim, Sheeny Vo, Nicholas Sweeney, Marie-Amandine Bonte, Ryan Rutherford, Dominic L Julian, Meixia Pan, Jacob Marsh, Andrea R Argouarch, Min Wu, Douglas W Scharre, Erica H Bell, Lawrence S Honig, Jean Paul Vonsattel, Geidy E Serrano, Thomas G Beach, Celeste M Karch, Aimee W Kao, Mark E Hester, Xianlin Han, and Hongjun Fu. GRAMD1B is a regulator of lipid homeostasis, autophagic flux and phosphorylated tau. Nat. Commun., 16(1):3312, April 2025. doi: 10.1038/s41467-025-58585-w.

[46] Jordan A Bairos, Uche Njoku, Maria Zafar, May G Akl, Lei Li, Gunes Parlakgul, Ana Paula Arruda, and Scott B Widenmaier. Sterol o-acyltransferase (SOAT/ACAT) activity is required to form cholesterol crystals in hepatocyte lipid droplets. Biochim. Biophys. Acta Mol. Cell Biol. Lipids, 1869(6):159512, August 2024. doi: 10.1016/j.bbalip.2024.159512.

[47] Norihito Shibata, Aaron F Carlin, Nathanael J Spann, Kaoru Saijo, Christopher S Morello, Jeffrey G McDonald, Casey E Romanoski, Mano R Maurya, Minna U Kaikkonen, Michael T Lam, Andrea Crotti, Donna Reichart, Jesse N Fox, Oswald Quehenberger, Christian R H Raetz, M Cameron Sullards, Robert C Murphy, Alfred H Merrill, Jr, H Alex Brown, Edward A Dennis, Eoin Fahy, Shankar Subramaniam, Douglas R Cavener, Deborah H Spector, David W Russell, and Christopher K Glass. 25-hydroxycholesterol activates the integrated stress response to reprogram transcription and translation in macrophages. J. Biol. Chem., 288(50):35812–35823, December 2013. doi: 10.1074/jbc.M113.519637.

[48] Puja Khanna, Joan Shuying Lee, Amornpun Sereemaspun, Haeryun Lee, and Gyeong Hun Baeg. GRAMD1B regulates cell migration in breast cancer cells through JAK/STAT and akt signaling. Sci. Rep., 8(1):9511, June 2018. doi: 10.1038/s41598-018-27864-6.

[49] Yifu Liu, Shengqiang Fu, Zhicheng Zhang, Siyuan Wang, Xiaofeng Cheng, Zhilong Li, Yi Ding, Ting Sun, and Ming Ma. GRAMD1A is a biomarker of kidney renal clear cell carcinoma and is associated with immune infiltration in the tumour microenvironment. Dis. Markers, 2022:5939021, July 2022. doi: 10.1155/2022/5939021.

[50] Seung-Soon Im and Timothy F Osborne. Liver x receptors in atherosclerosis and inflammation. Circ. Res., 108(8):996–1001, April 2011. doi: 10.1161/CIRCRESAHA.110.226878.

[51] Bruno Mesmin, Joëlle Bigay, Joachim Moser von Filseck, Sandra Lacas-Gervais, Guillaume Drin, and Bruno Antonny. A four-step cycle driven by PI(4)P hydrolysis directs sterol/PI(4)P exchange by the ER-Golgi tether OSBP. Cell, 155(4):830–843, November 2013. doi: 10.1016/j.cell.2013.09.056.

[52] Céline S C Hardy, Adam D Wegman, Mitchell J Waldran, Gary C Chan, and Adam T Waickman. Conventional and antibody-enhanced DENV infection of human macrophages induces differential immunotranscriptomic profiles. J. Virol., 99(3):e0196224, March 2025. doi: 10.1128/jvi.01962-24.

[53] Ramya Boddepalli, Harsh Chhajer, and Rahul Roy. Integrative Modelling of Innate Immune Response Dynamics during Virus Infection. *Preprint at bioRxiv*, June 2025. doi: 10.1101/2025.06.17.660089. URL http://biorxiv.org/lookup/doi/10.1101/2025.06.17.660089.

[54] Zeenat A. Shyr, Yu-Shan Cheng, Donald C. Lo, and Wei Zheng. Drug combination therapy for emerging viral diseases. Drug Discovery Today, 26(10):2367–2376, October 2021. ISSN 1878-5832. doi: 10.1016/j.drudis.2021.05.008.

[55] Rohan Narayan, Mansi Sharma, Rajesh Yadav, Abhijith Biji, Oyahida Khatun, Sumandeep Kaur, Aditi Kanojia, Christy Margrat Joy, Raju Rajmani, Pallavi Raj Sharma, Sharumathi Jeyasankar, Priya Rani, Radha Krishan Shandil, Shridhar Narayanan, Durga Chilakalapudi Rao, Vijaya Satchidanandam, Saumitra Das, Rachit Agarwal, and Shashank Tripathi. Picolinic acid is a broad-spectrum inhibitor of enveloped virus entry that restricts SARS-CoV-2 and influenza A virus in vivo. Cell Reports Medicine, 4(8):101127, August 2023. ISSN 26663791. doi: 10.1016/j.xcrm.2023.101127. URL https://linkinghub.elsevier.com/retrieve/pii/S2666379123002550.

[56] Kleber Juvenal Silva Farias, Paula Renata Lima Machado, Renato Ferreira de Almeida Junior, Ana Alice de Aquino, and Benedito Antônio Lopes da Fonseca. Chloroquine interferes with dengue-2 virus replication in u937 cells. Microbiology and Immunology, 58:318–326, 2014. ISSN 13480421. doi: 10.1111/1348-0421.12154.

[57] Eva Konkolova, Milan Dejmek, Hubert Hřebabecký, Michal Šála, Jiří Böserle, Radim Nencka, and Evzen Boura. Remdesivir triphosphate can efficiently inhibit the RNA-dependent RNA polymerase from various flaviviruses. Antiviral Research, 182:104899, October 2020. ISSN 01663542. doi: 10.1016/j.antiviral.2020.104899. URL https://linkinghub.elsevier.com/retrieve/pii/S0166354220303132.

[58] Obdulio García-Nicolás, Philip V’kovski, Nathalie J. Vielle, Nadine Ebert, Roland Züst, Jasmine Portmann, Hanspeter Stalder, Véronique Gaschen, Gabrielle Vieyres, Michael Stoffel, Matthias Schweizer, Artur Summerfield, Olivier Engler, Thomas Pietschmann, Daniel Todt, Marco P. Alves, Volker Thiel, and Stephanie Pfaender. The small-compound inhibitor k22 displays broad antiviral activity against different members of the family flaviviridae and offers potential as a panviral inhibitor. Antimicrobial Agents and Chemotherapy, 62(11):10.1128/aac.01206–18, 2018. doi: 10.1128/aac.01206-18. URL https://journals.asm.org/doi/abs/10.1128/aac.01206-18.

[59] Martin J Vincent, Eric Bergeron, Suzanne Benjannet, Bobbie R Erickson, Pierre E Rollin, Thomas G Ksiazek, Nabil G Seidah, and Stuart T Nichol. Chloroquine is a potent inhibitor of SARS coronavirus infection and spread. Virology Journal, 2(1):69, December 2005. ISSN 1743-422X. doi: 10.1186/1743-422X-2-69. URL https://virologyj.biomedcentral.com/articles/ 10.1186/1743-422X-2-69.

[60] Selvin Noé Palacios-Rápalo, Luis Adrián De Jesús-González, Carlos Daniel Cordero-Rivera, Carlos Noe Farfan-Morales, Juan Fidel Osuna-Ramos, Gustavo Martínez-Mier, Judith Quistián- Galván, Armando Muñoz-Pérez, Víctor Bernal-Dolores, Rosa María Del Ángel, and José Manuel Reyes-Ruiz. Cholesterol-Rich Lipid Rafts as Platforms for SARS-CoV-2 Entry. Frontiers in Immunology, 12:796855, December 2021. ISSN 1664-3224. doi: 10.3389/fimmu.2021.796855. URL https://www.frontiersin.org/articles/10.3389/fimmu.2021.796855/full.

[61] Marco M. Domingues, Bárbara Gomes, Axel Hollmann, and Nuno C. Santos. 25-hydroxycholesterol effect on membrane structure and mechanical properties. International Journal of Molecular Sciences, 22(5):2574, March 2021. ISSN 1422-0067. doi: 10.3390/ijms22052574. URL 10.3390/ijms22052574.

[62] Rubén Soto-Acosta, Clemente Mosso, Margot Cervantes-Salazar, Henry Puerta-Guardo, Fernando Medina, Liliana Favari, Juan E. Ludert, and Rosa María Del Angel. The increase in cholesterol levels at early stages after dengue virus infection correlates with an augment in LDL particle uptake and HMG-CoA reductase activity. Virology, 442(2):132–147, August 2013. ISSN 00426822. doi: 10.1016/j.virol.2013.04.003. URL https://linkinghub.elsevier.com/ retrieve/pii/S0042682213001979.

[63] Harsh Chhajer, Vaseef A. Rizvi, and Rahul Roy. Life cycle process dependencies of positive-sense RNA viruses suggest strategies for inhibiting productive cellular infection. *Journal of the Royal Society*, Interface, 18(184):20210401, November 2021. ISSN 1742-5662. doi: 10.1098/rsif.2021.0401.

[64] Eliana G Acosta, Anil Kumar, and Ralf Bartenschlager. Revisiting dengue virus–host cell interaction. In Advances in Virus Research, Advances in virus research, pages 1–109. Elsevier, 2014.

[65] T Ojala, M Pietikainen, and T Maenpaa. Multiresolution gray-scale and rotation invariant texture classification with local binary patterns. IEEE Trans. Pattern Anal. Mach. Intell., 24(7):971–987, July 2002.

[66] Peter J Costianes and Joseph B Plock. Gray-level co-occurrence matrices as features in edge enhanced images. In 2010 IEEE 39th Applied Imagery Pattern Recognition Workshop (AIPR). IEEE, October 2010.

[67] Naira Elazab, Wael Gab Allah, and Mohammed Elmogy. Computer-aided diagnosis system for grading brain tumor using histopathology images based on color and texture features. BMC Med. Imaging, 24(1):177, July 2024.

[68] Babraham Bioinformatics - Trim Galore! URL https://www.bioinformatics.babraham.ac.uk/projects/trim_galore/.

[69] Babraham Bioinformatics - FastQC A Quality Control tool for High Throughput Sequence Data. URL https://www.bioinformatics.babraham.ac.uk/projects/fastqc/.

[70] Peter W. Harrison, M. Ridwan Amode, Olanrewaju Austine-Orimoloye, Andrey G. Azov, Matthieu Barba, If Barnes, Arne Becker, Ruth Bennett, Andrew Berry, Jyothish Bhai, Simarpreet Kaur Bhurji, Sanjay Boddu, Paulo R. Branco Lins, Lucy Brooks, Shashank Budhanuru Ramaraju, Lahcen I. Campbell, Manuel Carbajo Martinez, Mehrnaz Charkhchi, Kapeel Chougule, Alexander Cockburn, Claire Davidson, Nishadi H. De Silva, Kamalkumar Dodiya, Sarah Donaldson, Bilal El Houdaigui, Tamara El Naboulsi, Reham Fatima, Carlos Garcia Giron, Thiago Genez, Dionysios Grigoriadis, Gurpreet S. Ghattaoraya, Jose Gonzalez Martinez, Tatiana A. Gurbich, Matthew Hardy, Zoe Hollis, Thibaut Hourlier, Toby Hunt, Mike Kay, Vinay Kaykala, Tuan Le, Diana Lemos, Disha Lodha, Diego Marques-Coelho, Gareth Maslen, Gabriela Alejandra Merino, Louisse Paola Mirabueno, Aleena Mushtaq, Syed Nakib Hossain, Denye N. Ogeh, Manoj Pandian Sakthivel, Anne Parker, Malcolm Perry, Ivana Piližota, Daniel Poppleton, Irina Prosovetskaia, Shriya Raj, José G. Pérez-Silva, Ahamed Imran Abdul Salam, Shradha Saraf, Nuno Saraiva-Agostinho, Dan Sheppard, Swati Sinha, Botond Sipos, Vasily Sitnik, William Stark, Emily Steed, Marie-Marthe Suner, Likhitha Surapaneni, Kyösti Sutinen, Francesca Floriana Tricomi, David Urbina-Gómez, Andres Veidenberg, Thomas A. Walsh, Doreen Ware, Elizabeth Wass, Natalie L. Willhoft, Jamie Allen, Jorge Alvarez-Jarreta, Marc Chakiachvili, Bethany Flint, Stefano Giorgetti, Leanne Haggerty, Garth R. Ilsley, Jon Keatley, Jane E. Loveland, Benjamin Moore, Jonathan M. Mudge, Guy Naamati, John Tate, Stephen J. Trevanion, Andrea Winterbottom, Adam Frankish, Sarah E. Hunt, Fiona Cunningham, Sarah Dyer, Robert D. Finn, Fergal J. Martin, and Andrew D. Yates. Ensembl 2024. Nucleic Acids Research, 52(D1):D891–D899, January 2024. ISSN 1362-4962. doi: 10.1093/nar/gkad1049.

[71] Alexander Dobin, Carrie A. Davis, Felix Schlesinger, Jorg Drenkow, Chris Zaleski, Sonali Jha, Philippe Batut, Mark Chaisson, and Thomas R. Gingeras. STAR: ultrafast universal RNA-seq aligner. Bioinformatics, 29(1):15–21, January 2013. ISSN 1367-4811, 1367-4803. doi: 10.1093/bioinformatics/bts635. URL https://academic.oup.com/bioinformatics/ article/29/1/15/272537.

[72] Yang Liao, Gordon K. Smyth, and Wei Shi. featureCounts: an efficient general purpose program for assigning sequence reads to genomic features. Bioinformatics, 30(7):923–930, April 2014. ISSN 1367-4811, 1367-4803. doi: 10.1093/bioinformatics/btt656. URL https://academic.oup.com/bioinformatics/article/30/7/923/232889.

[73] Boris Muzellec, Maria Teleńczuk, Vincent Cabeli, and Mathieu Andreux. PyDESeq2: a python package for bulk RNA-seq differential expression analysis. Bioinformatics, 39(9):btad547, September 2023. ISSN 1367-4811. doi: 10.1093/bioinformatics/btad547. URL https://academic.oup.com/bioinformatics/article/doi/10.1093/bioinformatics/btad547/7260507.

[74] Paul D. Thomas, Dustin Ebert, Anushya Muruganujan, Tremayne Mushayahama, LaurentPhilippe Albou, and Huaiyu Mi. Panther: Making genomescale phylogenetics accessible to all. Protein Science, 31(1):8–22, January 2022. ISSN 0961-8368, 1469-896X. doi: 10.1002/pro.4218. URL https://onlinelibrary.wiley.com/doi/10.1002/pro.4218.

[75] The Gene Ontology Consortium, Suzi A Aleksander, James Balhoff, Seth Carbon, J Michael Cherry, Harold J Drabkin, Dustin Ebert, Marc Feuermann, Pascale Gaudet, Nomi L Harris, David P Hill, Raymond Lee, Huaiyu Mi, Sierra Moxon, Christopher J Mungall, Anushya Muruganugan, Tremayne Mushayahama, Paul W Sternberg, Paul D Thomas, Kimberly Van Auken, Jolene Ramsey, Deborah A Siegele, Rex L Chisholm, Petra Fey, Maria Cristina Aspromonte, Maria Victoria Nugnes, Federica Quaglia, Silvio Tosatto, Michelle Giglio, Suvarna Nadendla, Giulia Antonazzo, Helen Attrill, Gil Dos Santos, Steven Marygold, Victor Strelets, Christopher J Tabone, Jim Thurmond, Pinglei Zhou, Saadullah H Ahmed, Praoparn Asanitthong, Diana Luna Buitrago, Meltem N Erdol, Matthew C Gage, Mohamed Ali Kadhum, Kan Yan Chloe Li, Miao Long, Aleksandra Michalak, Angeline Pesala, Armalya Pritazahra, Shirin C C Saverimuttu, Renzhi Su, Kate E Thurlow, Ruth C Lovering, Colin Logie, Snezhana Oliferenko, Judith Blake, Karen Christie, Lori Corbani, Mary E Dolan, Harold J Drabkin, David P Hill, Li Ni, Dmitry Sitnikov, Cynthia Smith, Alayne Cuzick, James Seager, Laurel Cooper, Justin Elser, Pankaj Jaiswal, Parul Gupta, Pankaj Jaiswal, Sushma Naithani, Manuel Lera-Ramirez, Kim Rutherford, Valerie Wood, Jeffrey L De Pons, Melinda R Dwinell, G Thomas Hayman, Mary L Kaldunski, Anne E Kwitek, Stanley J F Laulederkind, Marek A Tutaj, Mahima Vedi, Shur-Jen Wang, Peter D’Eustachio, Lucila Aimo, Kristian Axelsen, Alan Bridge, Nevila Hyka-Nouspikel, Anne Morgat, Suzi A Aleksander, J Michael Cherry, Stacia R Engel, Kalpana Karra, Stuart R Miyasato, Robert S Nash, Marek S Skrzypek, Shuai Weng, Edith D Wong, Erika Bakker, Tanya Z Berardini, Leonore Reiser, Andrea Auchincloss, Kristian Axelsen, Ghislaine Argoud-Puy, Marie-Claude Blatter, Emmanuel Boutet, Lionel Breuza, Alan Bridge, Cristina Casals-Casas, Elisabeth Coudert, Anne Estreicher, Maria Livia Famiglietti, Marc Feuermann, Arnaud Gos, Nadine Gruaz-Gumowski, Chantal Hulo, Nevila Hyka-Nouspikel, Florence Jungo, Philippe Le Mercier, Damien Lieberherr, Patrick Masson, Anne Morgat, Ivo Pedruzzi, Lucille Pourcel, Sylvain Poux, Catherine Rivoire, Shyamala Sundaram, Alex Bateman, Emily Bowler-Barnett, Hema Bye-A-Jee, Paul Denny, Alexandr Ignatchenko, Rizwan Ishtiaq, Antonia Lock, Yvonne Lussi, Michele Magrane, Maria J Martin, Sandra Orchard, Pedro Raposo, Elena Speretta, Nidhi Tyagi, Kate Warner, Rossana Zaru, Alexander D Diehl, Raymond Lee, Juancarlos Chan, Stavros Diamantakis, Daniela Raciti, Magdalena Zarowiecki, Malcolm Fisher, Christina James-Zorn, Virgilio Ponferrada, Aaron Zorn, Sridhar Ramachandran, Leyla Ruzicka, Monte Westerfield, Suzi A Aleksander, James Balhoff, Seth Carbon, J Michael Cherry, Harold J Drabkin, Dustin Ebert, Marc Feuermann, Pascale Gaudet, Nomi L Harris, David P Hill, Raymond Lee, Huaiyu Mi, Sierra Moxon, Christopher J Mungall, Anushya Muruganugan, Tremayne Mushayahama, Paul W Sternberg, Paul D Thomas, Kimberly Van Auken, Jolene Ramsey, Deborah A Siegele, Rex L Chisholm, Petra Fey, Maria Cristina Aspromonte, Maria Victoria Nugnes, Federica Quaglia, Silvio Tosatto, Michelle Giglio, Suvarna Nadendla, Giulia Antonazzo, Helen Attrill, Gil Dos Santos, Steven Marygold, Victor Strelets, Christopher J Tabone, Jim Thurmond, Pinglei Zhou, Saadullah H Ahmed, Praoparn Asanitthong, Diana Luna Buitrago, Meltem N Erdol, Matthew C Gage, Mohamed Ali Kadhum, Kan Yan Chloe Li, Miao Long, Aleksandra Michalak, Angeline Pesala, Armalya Pritazahra, Shirin C C Saverimuttu, Renzhi Su, Kate E Thurlow, Ruth C Lovering, Colin Logie, Snezhana Oliferenko, Judith Blake, Karen Christie, Lori Corbani, Mary E Dolan, Harold J Drabkin, David P Hill, Li Ni, Dmitry Sitnikov, Cynthia Smith, Alayne Cuzick, James Seager, Laurel Cooper, Justin Elser, Pankaj Jaiswal, Parul Gupta, Pankaj Jaiswal, Sushma Naithani, Manuel Lera-Ramirez, Kim Rutherford, Valerie Wood, Jeffrey L De Pons, Melinda R Dwinell, G Thomas Hayman, Mary L Kaldunski, Anne E Kwitek, Stanley J F Laulederkind, Marek A Tutaj, Mahima Vedi, Shur-Jen Wang, Peter D’Eustachio, Lucila Aimo, Kristian Axelsen, Alan Bridge, Nevila Hyka-Nouspikel, Anne Morgat, Suzi A Aleksander, J Michael Cherry, Stacia R Engel, Kalpana Karra, Stuart R Miyasato, Robert S Nash, Marek S Skrzypek, Shuai Weng, Edith D Wong, Erika Bakker, Tanya Z Berardini, Leonore Reiser, Andrea Auchincloss, Kristian Axelsen, Ghislaine Argoud-Puy, Marie-Claude Blatter, Emmanuel Boutet, Lionel Breuza, Alan Bridge, Cristina Casals-Casas, Elisabeth Coudert, Anne Estreicher, Maria Livia Famiglietti, Marc Feuermann, Arnaud Gos, Nadine Gruaz-Gumowski, Chantal Hulo, Nevila Hyka-Nouspikel, Florence Jungo, Philippe Le Mercier, Damien Lieberherr, Patrick Masson, Anne Morgat, Ivo Pedruzzi, Lucille Pourcel, Sylvain Poux, Catherine Rivoire, Shyamala Sundaram, Alex Bateman, Emily Bowler-Barnett, Hema Bye-A-Jee, Paul Denny, Alexandr Ignatchenko, Rizwan Ishtiaq, Antonia Lock, Yvonne Lussi, Michele Magrane, Maria J Martin, Sandra Orchard, Pedro Raposo, Elena Speretta, Nidhi Tyagi, Kate Warner, Rossana Zaru, Alexander D Diehl, Raymond Lee, Juancarlos Chan, Stavros Diamantakis, Daniela Raciti, Magdalena Zarowiecki, Malcolm Fisher, Christina James-Zorn, Virgilio Ponferrada, Aaron Zorn, Sridhar Ramachandran, Leyla Ruzicka, and Monte Westerfield. The Gene Ontology knowledgebase in 2023. GENETICS, 224(1):iyad031, May 2023. ISSN 1943-2631. doi: 10.1093/genetics/iyad031. URL https://academic.oup.com/genetics/article/doi/10.1093/genetics/iyad031/7068118.

[76] Michael Ashburner, Catherine A. Ball, Judith A. Blake, David Botstein, Heather Butler, J. Michael Cherry, Allan P. Davis, Kara Dolinski, Selina S. Dwight, Janan T. Eppig, Midori A. Harris, David P. Hill, Laurie Issel-Tarver, Andrew Kasarskis, Suzanna Lewis, John C. Matese, Joel E. Richardson, Martin Ringwald, Gerald M. Rubin, and Gavin Sherlock. Gene Ontology: tool for the unification of biology. Nature Genetics, 25(1):25–29, May 2000. ISSN 1061-4036, 1546-1718. doi: 10.1038/75556. URL https://www.nature.com/articles/ng0500_25.

[77] Alexander Lachmann, Denis Torre, Alexandra B. Keenan, Kathleen M. Jagodnik, Hoyjin J. Lee, Lily Wang, Moshe C. Silverstein, and Avi Ma’ayan. Massive mining of publicly available RNA-seq data from human and mouse. Nature Communications, 9(1):1366, April 2018. ISSN 2041-1723. doi: 10.1038/s41467-018-03751-6. URL https://www.nature.com/articles/s41467-018-03751-6.

[78] Zhuoqing Fang, Xinyuan Liu, and Gary Peltz. Gseapy: a comprehensive package for performing gene set enrichment analysis in python. Bioinformatics, 39(1):btac757, 11 2022. ISSN 1367-4811. doi: 10.1093/bioinformatics/btac757. URL 10.1093/bioinformatics/btac757.

[79] Reactome. Reactome pathways gene set, 91, December 2024. URL https://reactome.org/download-data/.

[80] Achinta Sannigrahi, Souradeepa Ghosh, Supratim Pradhan, Pulak Jana, Junaid Jibran Jawed, Subrata Majumdar, Syamal Roy, Sanat Karmakar, Budhaditya Mukherjee, and Krishnananda Chattopadhyay. Leishmania protein KMP-11 modulates cholesterol transport and membrane fluidity to facilitate host cell invasion. EMBO Reports, 25(12):5561–5598, October 2024. ISSN 1469-3178. doi: 10.1038/s44319-024-00302-7. URL https://www.embopress.org/doi/full/10.1038/s44319-024-00302-7.

